# The colocatome as a spatial -omic reveals shared microenvironment features between tumour–stroma assembloids and human lung cancer

**DOI:** 10.1101/2023.09.11.557278

**Authors:** Gina Bouchard, Weiruo Zhang, Irene Li, Ilayda Ilerten, Asmita Bhattacharya, Yuanyuan Li, Winston Trope, Joseph B Shrager, Calvin Kuo, Lu Tian, Amato J Giaccia, Sylvia K Plevritis

## Abstract

Computational frameworks to quantify and compare microenvironment spatial features of in-vitro patient-derived models and clinical specimens are needed. Here, we acquired and analysed multiplexed immunofluorescence images of human lung adenocarcinoma (LUAD) alongside tumour– stroma assembloids constructed with organoids and fibroblasts harvested from the leading edge (Tumour-Adjacent Fibroblasts;TAFs) or core (Tumour Core Fibroblasts;TCFs) of human LUAD. We introduce the concept of the “colocatome” as a spatial -omic dimension to catalogue all proximate and distant colocalisations between malignant and fibroblast subpopulations in both the assembloids and clinical specimens. The colocatome expands upon the colocalisation quotient (CLQ) through a nomalisation strategy that involves permutation analysis and thereby allows comparisons of CLQs under different conditions. Using colocatome analysis, we report that both TAFs and TCFs protected cancer cells from targeted oncogene treatment by uniquely reorganising the tumour–stroma cytoarchitecture, rather than by promoting cellular heterogeneity or selection. Moreover, we show that the assembloids’ colocatome recapitulates the tumour–stroma cytoarchitecture defining the tumour microenvironment of LUAD clinical samples and thereby can serve as a functional spatial readout to guide translational discoveries.

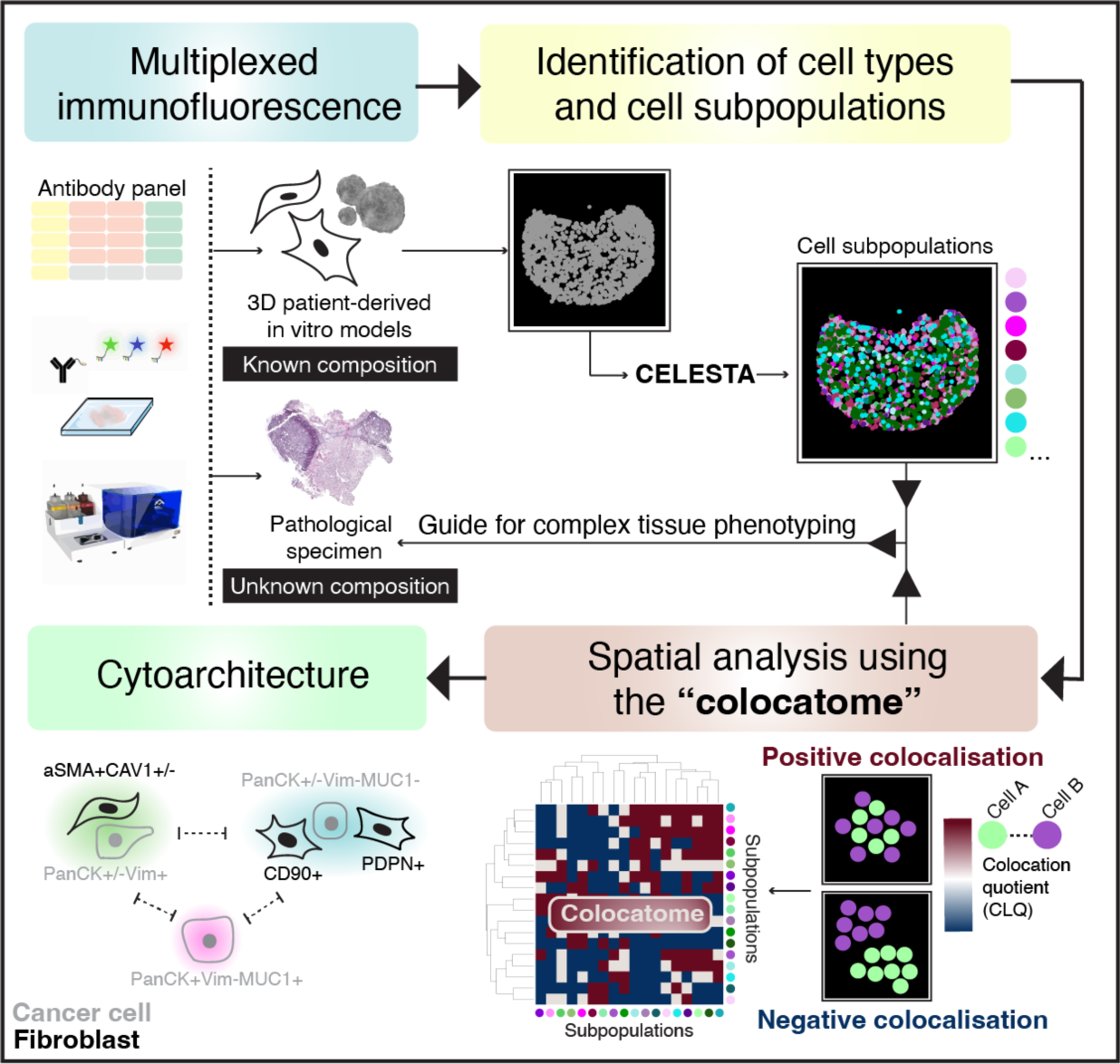

## Introduction

The emergence of highly multiplexed spatial -omics technologies is propelling translational oncology research forward by enabling a more comprehensive understanding of the cellular organisation within the tumour microenvironment (TME)^1^. In parallel, translational research using 3D patient-derived models is providing deeper insights into patient-specific TME functions by offering high throughput experimental platforms for probing cell–cell interactions underlying human pathobiology^2^. Given the rapid adoption of these technologies, computational approaches are urgently needed to quantitatively assess the spatial organisation of in vitro patient-derived models in a manner that is transferable to clinical specimens. For this purpose, the term “cytoarchitecture,” frequently employed to describe the structural arrangement of neurons^3^, is being adopted here to characterise the spatial co-occurrence of cell subpopulations in the TME. While numerous sudies have profiled tumour–stroma interactions using 3D patient-derived models^4–6^, here we focus on the spatial characterisation of the cytoarchitecture since the translational relevance of this information is largely unexplored.

We propose a quantitative approach to characterise the tumour–stroma cytoarchitecture of lung adenocarcinoma (LUAD) assembloids and clinical samples through use of the “colocatome”. We introduce the “colocatome” as an –omic dimension that provides a comprehensive spatial readout of the cytoarchitecture by cataloguing all positive (proximate) and negative (distant) cell–cell colocalisations between cell subpopulation pairs from distinct assembloid models in a manner that is transferable to clinical samples. The focus of this study is on the tumour–stroma colocatome of lung adenocarcinoma (LUAD) as lung cancer is the leading cause of cancer-related death in the world^8^, with LUAD being the main subtype of lung cancer; however, the colocatome concept is generalisable. To quantify the tumour–stroma LUAD colocatome, we generated assembloids^7^ using LUAD epithelial organoids and previously characterised Cancer-Associated Fibroblasts (CAFs) isolated from spatially distinct tumour sites (tumour edge vs tumour core)^9^, hereon referred to as regionally distinct CAFs. Using our colocatome analysis, we test the hypothesis that regionally distinct CAFs uniquely influence the tumour–stroma cytoarchitecture of the LUAD tumour– stroma assembloids. The rationale for this hypothesis stems from our previous work where we demonstrated that regionally distinct CAFs from the tumour edge, namely tumour-adjacent fibroblasts (TAFs), versus tumour core fibroblasts (TCFs) promoted striking morphological and transcriptomic differences when cocultured in 2D with cancer cells; moreover, we demonstrated the promigratory features of cancer cells cocultured with TAFs vs TCFs^9^.

To quantify and compare the cytoarchitecture of TAF–PDO and TCF–PDO assembloids, we acquired multiplexed immunofluorescence (mIF) images of assembloid sections using a PhenoCycler (formerly known as CODEX)^10^, on which we performed quantitative spatial biology analyses. To identify the two major cell subtypes (malignant and fibroblast) and their respective cell subpopulations within the assembloids, we extended the capabilities of CELESTA^11^, a semi-supervised machine learning algorithm for cell type identification. Then, we applied the colocation quotient (CLQ) to quantitatively characterise the spatial organisation of fibroblasts and cancer cell subpopulations^11,12^. We present the colocatome analysis as a method that expands upon the colocalisation quotient (CLQ) analysis by utlising a nomalisation strategy that involves spatial permutation testing and thereby allows CLQs to be compared across different conditions, including assembloid models and tissue samples.

Through our colocatome analysis of the tumour–stroma assembloids, we find that regionally distinct CAFs exhibit unique tumour–stroma cytoarchitecture and promote treatment resistance through distinct spatial reorganisation rather than specific cell subpopulation selection. Moreover, we show that the colocatome derived from our tumour–stroma LUAD assembloids recapitulates the tumour–stroma colocatome associated with specific LUAD histopathological growth patterns. Thus, the spatial features of assembloids can be transferred to clinical specimens and thereby help to guide complex tissues phenotyping. In summary, we underscore the power of coupling spatial –omics technologies with computational tools to elucidate the cytoarchitecture of 3D assembloids. Importantly, by transferring spatial features from assembloids to clinical samples, we demonstrate the utility of the “colocatome” as a new functional readout and –omic dimension to answer complex spatial biology question about the tumour microenvironment and beyond.

## Results

### Regionally distinct CAFs exhibit unique spatial organisation in tumour–stroma assembloid models

To model tumour–stroma heterogeneity, we established CAF–PDO assembloids and investigated whether regionally distinct CAF originating from the tumour leading edge (TAFs) vs core (TCFs) influence cell composition and organisation. Assembloids were generated from single-cell suspensions of two cell types (cancer cells and fibroblasts), incubated for up to 10 days, embedded, sectioned, and stained with antibodies using immunofluorescence (IF), as if they were pathological specimens (Fig. 1a). IF results showed that both assembloid models displayed inter- (TAFs vs TCFs) and intra- (periphery vs centre) spatial heterogeneity. TAF–PDO assembloids were enriched in periphery of the matrix dome, and displayed lower cell density in the centre (Fig. 1a). In contrast, TCF–PDO assembloids were more homogeneously organised within the matrix dome. Our IF results suggest that fibroblasts impact the spatial organisation of cancer cells depending on their origin in the tumour; we find this finding to be statistically significant by combining mIF with a quantitative spatial analysis approach, as described next and shown in the study workflow (Fig. 1b).

**Fig. 1.**
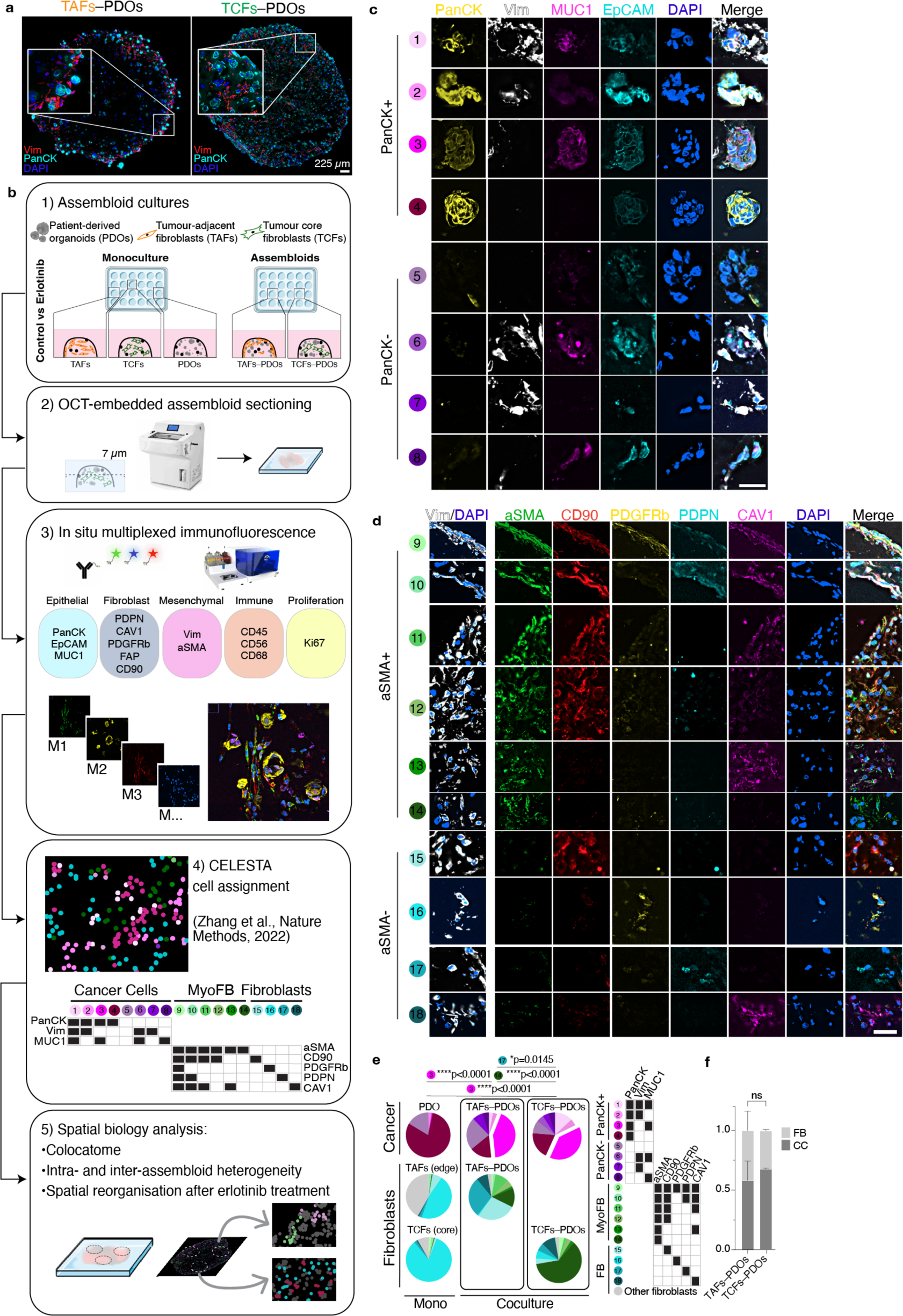
Cell composition of TAF–PDO and TCF–PDO LUAD assembloids. **a**, Traditional IF images with two markers (PacnCK, Vim) and one nuclei stain (DAPI) of OCT sections of TAF–PDO (left) and TCF–PDO (right) LUAD assembloids. **b**, Study workflow and mIF analysis pipeline. **c**, Representative PhenoCycler images of assembloids showing cancer cell and (**d**) fibroblast subpopulations identified by CELESTA. Scale bar, 30 µm. **e**,**f,** Representative example of subpopulation proportions in TAF–PDO and TCF–PDO LUAD assembloids. Statistical significance was determined using two-way ANOVA and Fisher’s least significant difference between subpopulations and assembloid conditions. Error bars indicate standard deviation of the mean. Additional biological replicate can be found in Extended data Fig. 1a. Assembloids were generated with matched fibroblasts from the same patient and data represent biological replicates from two independent experiments. LUAD, lung adenocarcinoma; CC, cancer cell; FB, fibroblast. OCT, optimum cutting temperature; Vim, vimentin; PanCK, pan-cytokeratin; EpCAM, epithelial cell adhesion molecule; MUC1, mucin 1; PDPN, podoplanin; CAV1, caveolin 1; PDGFRb, platelet derived growth factor receptor beta; FAP, fibroblast activation protein; CD, cluster of differentiation; aSMA, smooth muscle actin alpha 2.

### TAF–PDO vs TCF–PDO assembloids reveal distinct fibroblast subpopulations but similar cancer cell subpopulations

To conduct a more comprehensive quantitative assessment of the assembloids’ spatial organisation, we used the mIF PhenoCycler technology from Akoya Biosciences. We acquired images from a panel of 15 markers, mainly constituted of epithelial and fibroblast markers, to which we applied our computational pipeline to resolve the cytoarchitecture. A early step in this pipeline is to identify cell subpopulations at single cell resolution. After cell segmentation, a common approach for identifying cell subpopulations involves clustering cells with similar expression patterns. However, the assignment of clusters is subjective, time-consuming and requires manual assessment. Moreover, some clusters of mixed cell subpopulations can be difficult to annotate. To overcome these challenges, we expanded the capabilities of the CELESTA algorithm. CELESTA is a semi-supervised machine learning tool that does not involve manual gating or clustering, but uses prior or partial knowledge of the cell expression profile and spatial information^11^. CELESTA allowed us to identify the main cell subtypes in the assembloids (cancer cells and fibroblasts), as well as numerous subpopulations.

Identifying precise subpopulations in complex tissues of unknown composition is difficult to achieve, even with the help of an expert pathologist. To address this challenge, we used pre-defined fibroblasts and organoid cultures of known characteristics to build tumour–stroma assembloids, with the goal of using the subpopulations identified in the assembloids as a guide to identifying similar subpopulations in clinical samples. For subpopulation assignment in the assembloids, we leveraged the customable thresholds and rapid iteration features of CELESTA, which allowed us to rapidly test every combination of fibroblast and epithelial markers (Extended Data Fig. 1a). In total, we identified 18 subpopulations in the assembloids (Fig. 1c–f): 8 cancer cells and 10 fibroblasts, with additional fibroblast subpopulations specific to monocultures (Extended Data Fig. 1b and Supplementary Table 1). The subpopulations were segregated between PanCK+ and PanCK- for cancer cells, αSMA+ (myofibroblasts) and αSMA- for fibroblasts, and validated on the original images. Cancer cells displayed broad morphological differences, ranging from adherent to mesenchymal cell bundles (Fig. 1c), and myofibroblasts appeared more elongated than αSMA-fibroblasts (Fig. 1d). We found that PDO 3D monocultures were largely composed of PanCK+Vim-MUC1-cancer cells (subpopulation #4), whereas in the presence of TAFs and TCFs, the dominant cancer cell subpopulation was PanCK+Vim-MUC1+ (subpopulation #3, p<0.0001, p<0.0001). As for fibroblasts, cancer cells stimulated their transformation into myofibroblasts (mainly absent from monocultures). Even though TAF–PDO assembloids were enriched for PDPN+ fibroblasts (subpopulation #17, p=0.0145) and TCF–PDO assembloids were mainly constituted of aSMA+ fibroblasts (subpopulation #14, p<0.0001) (Fig. 1e), the assembloids exhibited similar cancer cell heterogeneity. Interestingly, the initial 1:1 cancer cell:fibroblast ratio was mostly conserved in both assembloids, and neither TAFs or TCFs significantly induced cancer cell proliferation (Fig. 1f). Next, we quantified whether certain cell subpopulations were enriched in either the periphery or the centre of the assembloids. TAFs subpopulations #9-11 (expressing three or more fibroblast markers) were significantly enriched at the periphery (p<0.0001) and subpopulations #14-18 (expressing 1 marker) at the centre (p=0.0036, Extended Data Fig. 1c,d). In contrast, TCFs and cancer cell subpopulations were not differentially enriched in either zone (Extended Data Fig. 1d and Extended Data Fig. 2-5).

### Colocatome analysis demonstrates that TAFs–PDOs vs TCFs–PDOs exhibit distinct tumour–stroma cytoarchitecture

To quantitatively evaluate and compare the cytoarchitecture of the TAFs–PDO and TCFs– PDO assembloid models, we introduce the general concept of the colocatome as a spatial -omic dimension. In the context of tumour–stroma assembloids, the colocatome catalogues all positive (proximate) and negative (distant) cell–cell colocalisations between cancer cell and fibroblast subpopulations. The colocatome expands upon the colocalisation quotient (CLQ), a measure designed to assess the spatial association between different categories within a population that may exhibit spatial organisation^11,12^. To construct the colocatome, we first compute the CLQ value for each pair of cell subpopulations, using the 20 nearest neighbors for each index cell subpopulaton. We then examine which cell subpopulation pairs are likely to be colocated (proximate), as indicated by a high CLQ value, or spatially distant, as indicated by a low CLQ value (Fig. 2a). As detailed in the Methods, we expand on prior work by assessing the significance of the CLQ values by randomly permuting the cell subpopulation label of each cell while preserving the cell subpopulation composition to generate the CLQ null distribution. Observed CLQ values falling within vs outside of the tail of their respective null distribution were considered statistically significant (Fig. 2b, Supplementary Table 2). This permutation approach takes into account the prevalence of cell subpopulations when determining significance. For example, when the CLQ is computed among two highly prevalent cell subpopulations, the null CLQ distribution tends to narrow, while rare subpopulations lead to broader null CLQ distributions. In the latter case, the observed CLQ values are more likely to be located within the null distribution and thereby considered non-significant. We then used the information from the null distribution to normalise the CLQs and generate the colocatome, which allows us to compare spatial features (positive and negative colocalisations included) between cell subpopulation pairs across different conditions (i.e., TAFs vs TCF assembloid), and samples types (i.e., pathological specimens vs. assembloids).

**Fig. 2.**
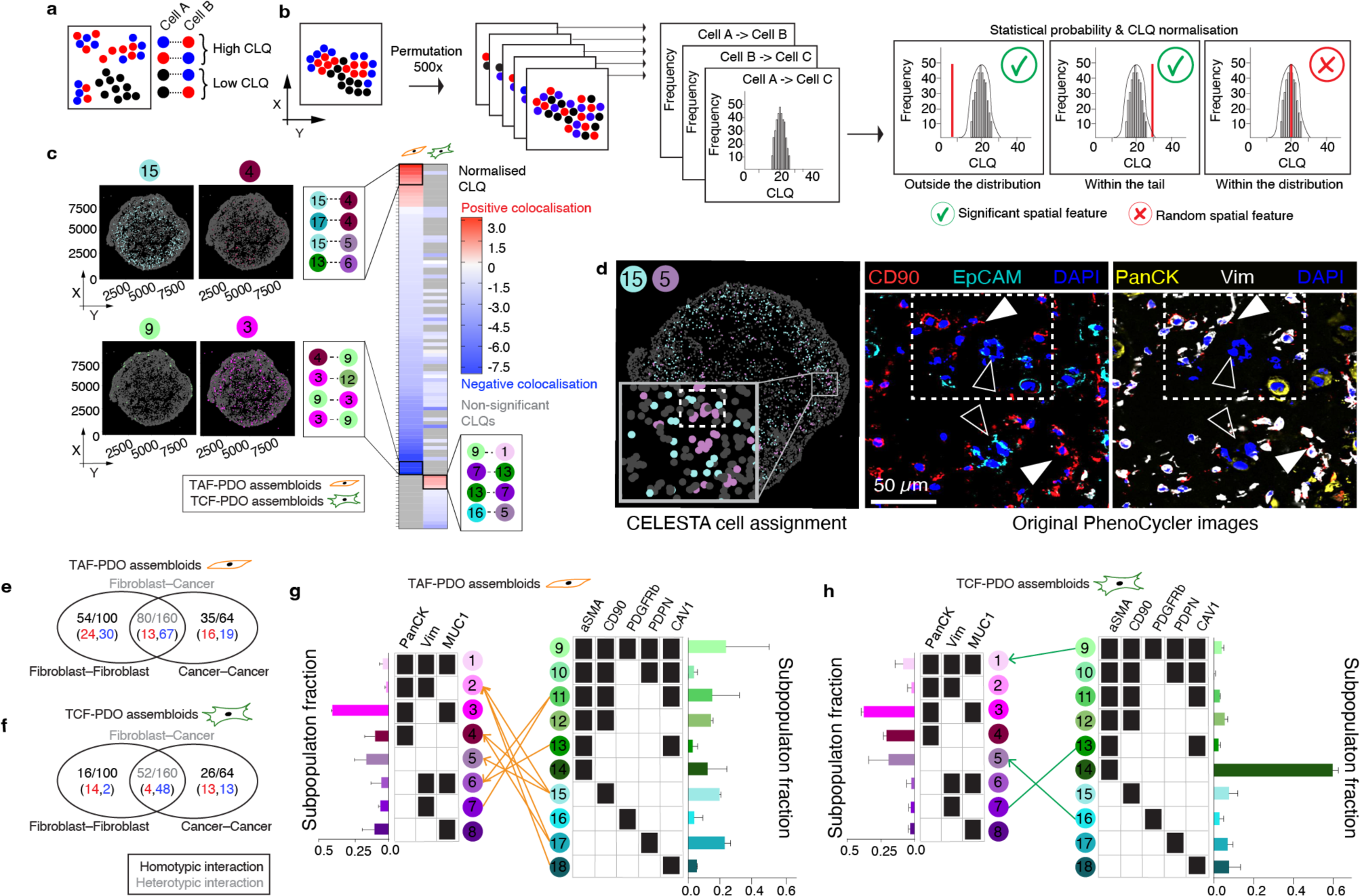
Colocatome analysis enables a comparisons of cell–cell colocalisations in TAF–PDO versus TCF–PDO assembloids. **a**, Schematic representation of the colocation quotient (CLQ) and (**b**) workflow of the permutation analysis to determine statistically significant cell–cell colocalisations. **c**, Heatmaps showing statistically significant heterotypic negative (blue) and positive colocalisations (red) in TAF–PDO and TCF–PDO assembloids with example plots showing the subpopulation fractions. CLQs and their corresponding p values can be found in Supplementary Table 2. **d**, Validation of a significant colocalisation on original PhenoCycler images. White arrowheads indicate fibroblasts and empty arrowheads indicate cancer cells. Venn diagram showing the number of homotypic and heterotypic interections in TAF–PDO **(e)** and TCF–PDO **(f)** assembloids. Red vs blue numbers indicate positive vs negative colocalisations, respectively. Bipartite graph showing the heterotypic colocalisations of proximal partners found in TAF **(g)** and TCF **(h)** assembloids with corresponding bar graphs showing the subpopulation fractions. Lines without arrowheads represent bidirectional colocaliscations, and lines with arrowheads represent unidirectional colocalisations.

Based on the colocatome analysis, we quantified statitically significant positive and negative homotypic (cancer–cancer or fibroblast–fibroblast) and heterotypic (cancer–fibroblast) colocalisations, in both assembloid systems. Our colocatome analysis revealed that TAF–PDO assembloids exhibited a more organised cytoarchitecture, evidenced with approximately twice as many significant spatial features compared to TCF–PDO assembloids (Fig. 2c-h). Surprisingly, over 80% of the total heterotypic spatial features observed in both assembloid subtypes were negatively colocalised, suggesting that cancer cells and fibroblasts are primarily separated from each other rather than being intermixed proximate neighbours (Fig. 2c). Interestingly, no positive cancer–fibroblast colocalisations were shared between the TAF–PDO and TCF–PDO assembloids and, cell subpopulations with higher cell counts did not necessarily exhibit a higher number of statistically significant proximal neighbors. For instance, the largest cancer cell subpopulation (#3: PanCK+Vim-MUC1+) did not positively colocalise with any fibroblast (Fig. 2g,h). In summary, these results demonstrate that regionally distinct fibroblasts contribute to unique tumour–stroma assembloid cytoarchitecture in the tumour–stroma assembloids.

### Colocatome analysis shows that erlotinib reorganises the tumour–stroma assembloids’ cytoarchitecture without selecting for cell subpopulations

Little is known about the effects of drugs on tissue cytoarchitecture. CAFs have been shown to protect cancer cells through various mechanisms and have been proposed as a therapeutic target in cancer^13,14^. Therefore, we asked the question whether fibroblasts can induce drug resistance through cytoarchitectural rearrangement. We utilised erlotinib, which targets the epidermal growth factor receptor (EGFR), as a means to disrupt the cytoarchitecture of the assembloids. Given that the LUAD PDOs used in this study are EGFR+ (Supplementary Table 3), we found that erlotinib reduced cell density of the PDO monocultures (p=0.04, (Extended data Fig. 6 and 7). Interestingly, no significant decrease of cell density was measured for either erlotinib-treated assembloids, compared to their respective treatment-naïve controls, indicating that both TAFs and TCFs can protect cancer cells from erlotinib-induced cell death (Fig. 3a).

**Fig. 3.**
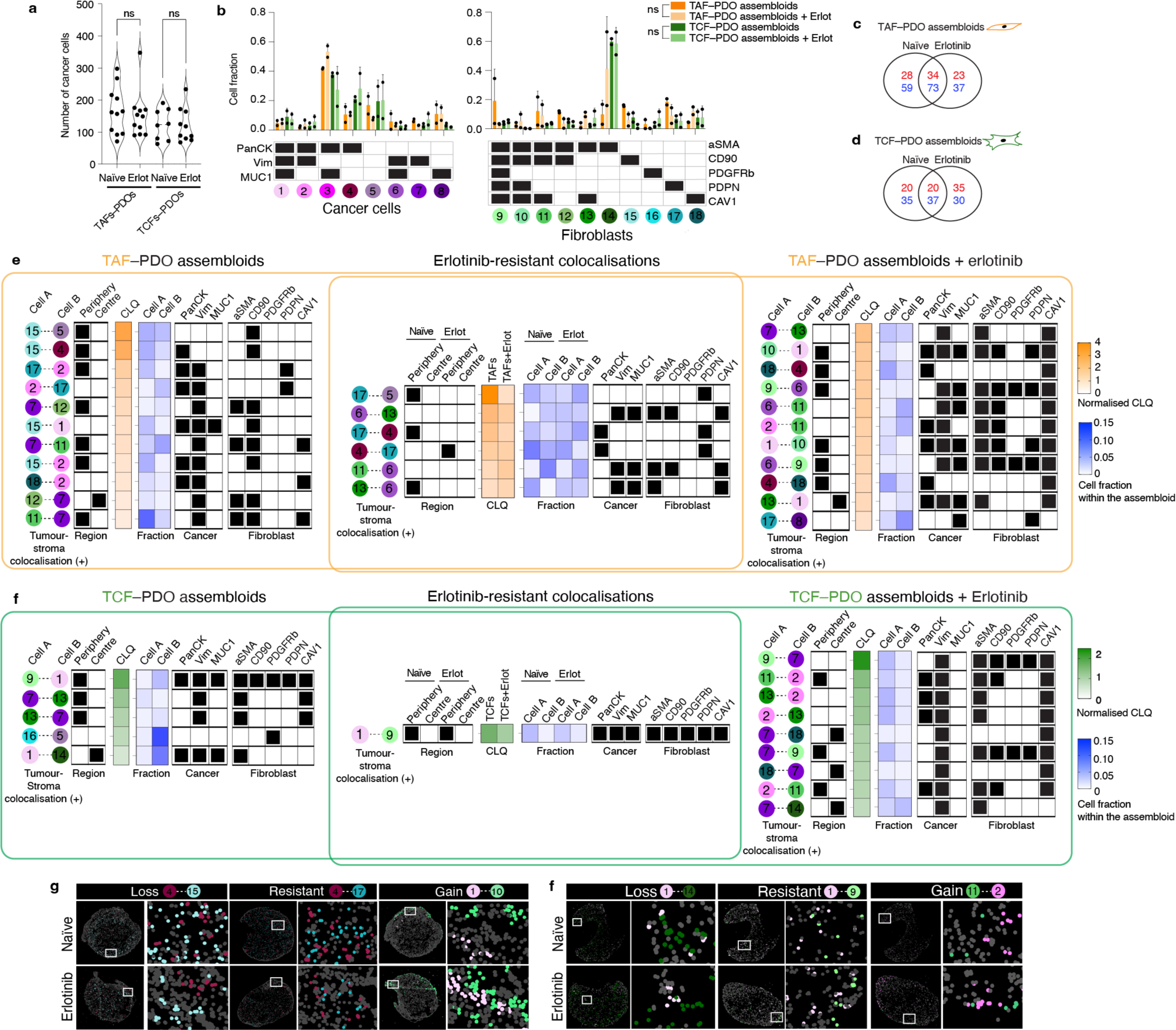
Effect of erlotinib treatment on tumour–stroma assembloids. **a**, Violin plot representing cancer cell density in treatment-naïve and erlotinib-treated assembloids. Quantification of 7 to 12 representative areas (depending on the size of the assembloid section) per condition. Statistical significance was determined using one-way ANOVA. Biological replicate can be found at Extended Data Fig. 8a. **b**, Bar graphs showing the fraction of each cell subpopulation before and after erlotinib from two independent biological replicate per condition. Error bars indicate standard deviation of the mean. Statistical significance was determined using two-way ANOVA. Venn diagram showing the number of positive (red) and negative colocalisations (blue) in TAF–PDO **(c)** and TCF–PDO **(d)** assembloids. **e**,**f** Heatmaps representing normalised CLQ values of statistically significant heterotypic positive colocalisations only and subpopulation fractions in treatment-naïve and and erlotinib-treated assembloids. The normalised CLQs and their corresponding p-values can be found in the Supplementary Table 4. **g,** CELESTA cell assignment plots showing representative examples of gains, losses and resistant heterotypic colocalisations.

Strikingly, none of the cell subpopulations were significantly enriched or depleted after treatment in either assembloid model (Fig. 3b). Instead, erlotinib induced numerous gains and losses of colocalisations in both TAF–PDO and TCF–PDO assembloids, with many of these changes being enriched either at the periphery or centre of the assembloids (Fig. 3c–f and Extended data Fig. 8). Numerous cancer–fibroblast positive colocalisations, particularly in TAF–PDO assembloids, persisted under treatment (Fig. 3e) and a significant number of new positive colocalisations emerged after erlotinib treatment in both systems, predominantly involving Vim+ cancer cells and CAV1+ or αSMA+CAV1+ fibroblasts (Fig. 3e,f). Although erlotinib induced a partial epithelial-mesenchymal transition (EMT) in the PDO monocultures after 72 hours, which was associated with elevated levels of O-GlcNAc, MDR1, and MUC1, known mechanisms of erlotinib resistance^15–17^, this effect was modest (Extended data Fig. 6 and 7) and unlikely to have influenced the cytoarchitectural reorganisation. Overall, our findings indicate that both TAFs and TCFs can protect cancer cells from erlotinib, primarily through unique reorganisation of the tumour–stroma spatial cytoarchitecture rather than promoting heterogeneity or cell subpopulation selection.

### Colocatome analysis enables discovery of treatment-resistant tumour–stroma cytoarchitecture

To resolve the cytoarchitecture of the assembloids across treatment-naïve and erlotinib-treated conditions, we grouped all statistically significant cell–cell colocalisations existing under at least one condition into a ternary matrix of three possibilities. Briefly, negative colocalisations were assigned to −1, positive colocalisations were assigned to 1 and unsignificant spatial features were assigned to 0; this allowed us to establish a comprehensive and simplified reference of spatial features that we designate as the composite tumour– stroma assembloid colocatome. Among other things, this composite colocatome revealed that homotypic colocalisations were enriched compared to heterotypic colocalisations (Fig. 4a).

**Fig. 4.**
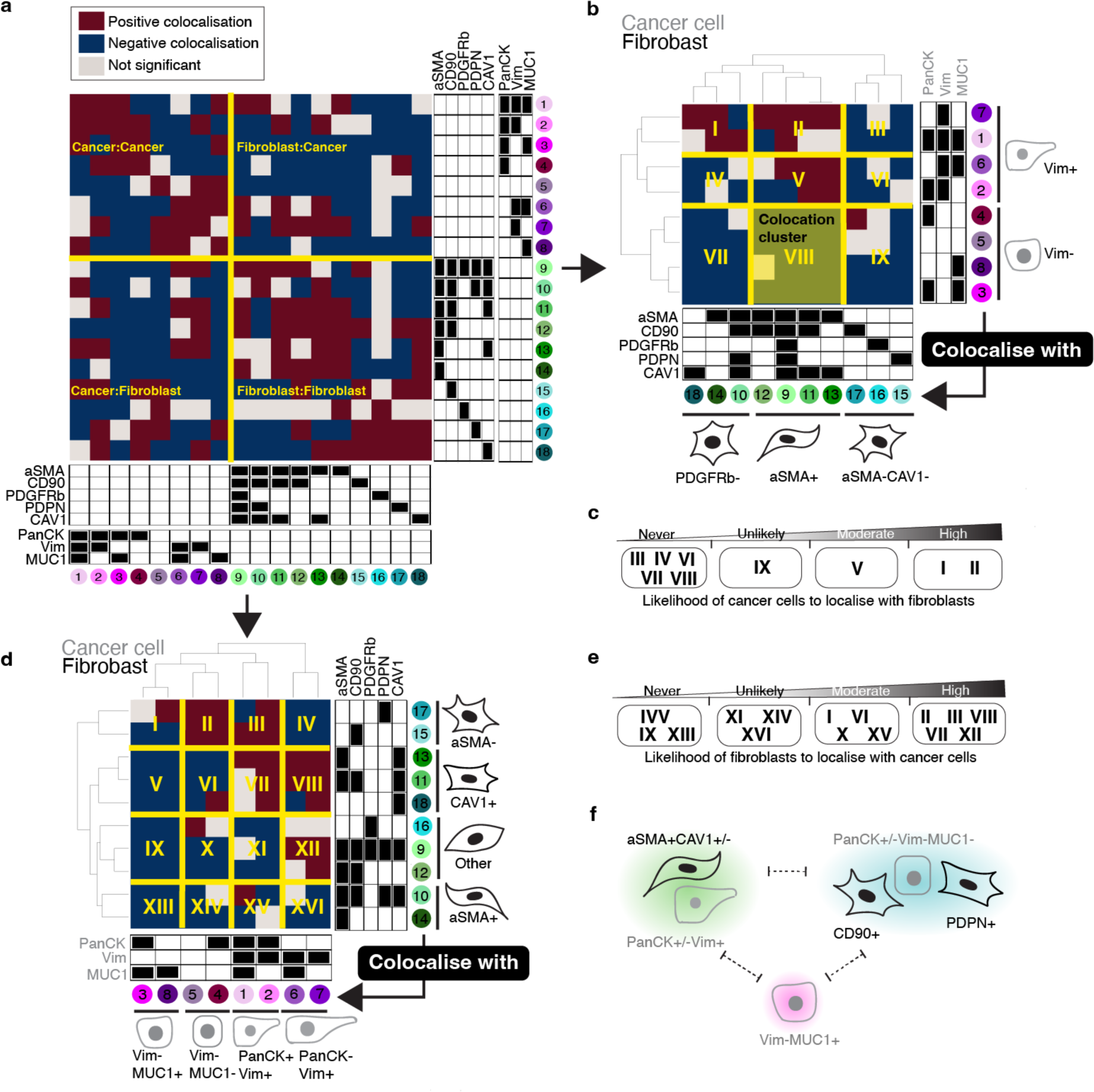
The tumour–stroma colocatome. **a**, Heatmap of the colocalisation possibilities encompassing all assembloid conditions generated in this study to establish a reference tumour–stroma colocatome. **b-e,** Hierarchical clustering of the tumour–stroma colocatome. Colocation clusters highlight groups of cells that colocalise with shared partners and indicate the likelihood of cancer cells to colocalise with fibroblasts or vice-versa. **f,** cartoon summary of the cancer–fibroblast cytoarchitecture inferred from the colocatome. Dotted lines indicate segregated spatial features that do not intermix.

Using hierarchical clustering on the heterotypic composite colocatome, we identified distinct colocation clusters (see roman numerals, Fig. 4b-e). In particular, these clusters highlight groups of cancer–fibroblast pairs that spatially behave similarly. Prior work has shown that fibroblasts promote EMT in cancer cells, and cancer cells sustain the myofibroblast phenotype in fibroblasts^18–20^. We corroborated these results by showing that Vim(-) cancer cells are unlikely to colocalise with myofibroblasts (aSMA+) (Fig. 4b-f), while myofibroblasts are generally found in proximity of Vim(+) cancer cells. In addition to confirming established cancer–fibroblast colocalisations, our composite colocatome analysis provided insights into previously unreported spatial configurations within the TME. Notably, we observed that MUC1+ cancer cells are unlikely to be found in proximity with fibroblasts. For instance, the cancer cell subpopulation #3 (PanCK+Vim-MUC1+) is not found in proximity with any fibroblast subtype regardless of the treatment condition. Moreover, we find that CD90+ fibroblasts are regularly found in the vicinity of cancer cells, while the inverse association is not observed, showing that spatial relationships between cancer cells and fibroblasts are asymetrical (Fig. 4b-e). Overall, the composite colocatome inferred from tumour–stroma assembloids validated known cancer–fibroblast colocalisations and revealed new cytoarchitectural features.

### Colocatome analysis demonstrates that colocalisations derived from tumour–stroma assembloids are recapitulated in LUAD clinical samples

To assess the clinical relevance of the colocatome, we generated the colocatome from histologic growth patterns derived from utilised whole slide mIF images LUAD clinical samples using the assembloid colocatome as a guide. An expert pathologist delineated 13 histological regions, assigned as either lepidic, acinar, or solid. Of note, the predominance of these histologic growth patterns have prognostic significance: lepidic predominant tumours are associated with a good prognosis due to their noninvasive nature, whereas acinar and solid predominant tumours are associated with worse prognosis, with solid worse than acinar^21,22^. Out of all the spatial features identified across the 13 histological regions, most (69/75) were observed in the composite colocatome generated from the assembloids (Fig. 5a-c, Supplementary Table 5). Hierarchical clustering based on the top 25% most variable colocalisations among the histologic growth patterns showed that solid regions mostly grouped with the TAF–PDO assembloids (green cluster 4, Fig. 5d), whereas the lepidic and acinar regions grouped predominantly with TCF–PDO assembloids (pink cluster 3, Fig. 5d). These results reinforce our published findings regarding the more tumour-promoting role of TAFs versus TCFs^23^ by associating TAF–PDOs’ spatial features with the worse prognostic growth patterns (solid) and TCF–PDOs’ spatial features with the better prognostic growth patterns (lepidic and acinar). To further demonstrate the relevance of the colocatome, we performed hierarchical clustering based on cell composition only. No clear association was found between the assembloids and the growth patterns, revealing the added value of this -omic dimension over analysis of cell composition only (Fig. 5e). Of note, additional information can be gleaned from the hierarchical analysis of the colocalisations: acinar regions from different clusters displayed distinct cytoarchitecture suggesting a previously undescribed spatial heterogeneity among acinar regions (Extended data Fig. 11).

**Fig. 5.**
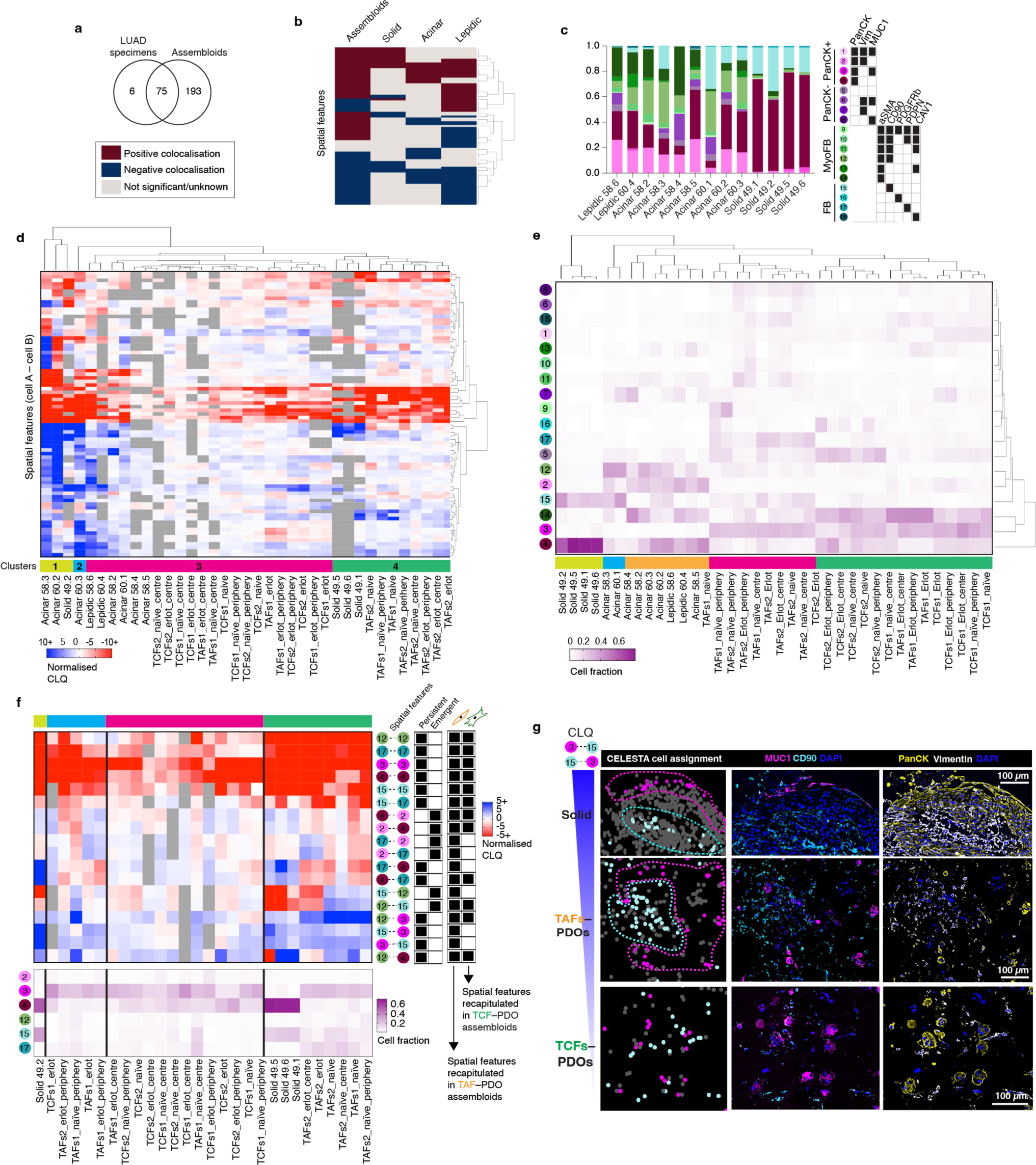
The colocatome from tumour–stroma recapitulate LUAD spatial features. **a**, Venn Diagram and heatmap (**b**) showing the number of significant anti- and colocalisations found in the LUAD specimens and assembloids. **c**, Cell proportions of each histopathological regions identified with CELESTA. **d**, Heatmap showing the normalised CLQ of the top 25% variable spatial features between clinical specimens. All p values can be found in Supplementary Table 5. **e,** Hierarchical clustering of the LUAD specimens and assembloids according to their cell composition. **f,** Heatmap of the spatial features identified from the LUAD solid regions integrated with the erlotinib-resistant spatial features inferred from the reference colocatome. **g,** Representative example of a negative colocalisation from a solid region recapitulated in the TAF–PDO, but not TCF–PDO assembloid in vitro model.

To identify potential spatial features associated with treatment resistance in clinical specimens, we next compared the statistically significant persistant and emergent colocalisations identified in the treatment-naïve vs erlotinib-treated assembloids that overlap with each of the histological growth patterns. We found that erlotinib-resistant spatial features from solid growth patterns were predominantly recapitulated in TAF–PDO assembloids (Fig. 5f). This finding further supports our previous finding by demonstrating a more pro-tumour progression role of TAFs compared to TCFs, and supports the use of TAF–PDO assembloids as more suitable than the TCF–PDO assembloids to recapitulate solid spatial features associated with treatment resistance. To further show how TAF–PDO assembloids may accurately replicate LUAD solid spatial features, we highlight a cancer–fibroblast colocalisation that was reproduced exclusively in TAF–PDO assembloids. Specifically, PanCK+Vim-MUC1+ cancer cells (#3) negatively colocalise with CD90+ fibroblasts (#15), which appeared as segregated compartments in LUAD specimens and TAF–PDO assembloids, but not in the the TCF–PDO assembloids (Fig. 5g). Similarly, we identified that acinar-like colocalisation may be more accurately recapitulated in TCF–PDO assembloids. Using hierachical clustering, we observed that resistant acinar-like spatial features were influenced by whether the colocalisation is located in the centre or at the periphery of the assembloids, which we did not oberve for TAF–PDO assembloids (Extended Data Fig. 13 and 14). Overall, these findings suggest that using comparative colocatome analysis with different assembloids alongside clinical specimens can help identify spatial biomarkers with clinical relevance and suitable validation platforms for subsequent functional analysis

## Discussion

In this study, we present the concept of the colocatome as a spatial -omic to characterise and compare the cytoarchitecture of in vitro models among themselves and with tissue samples. We apply the colocatome to study the tumour–stroma cytoarchitecture using TAF–PDO and TCF–PDO assembloids, as well as LUAD clinical specimens. Overall, our findings support comparative colocatome analysis between assembloid models and clinical specimens as means to guide the identification of clinically-relevant spatial features in the assembloid models for downstream functional analysis with translational relevance.

When comparing the colocatome of TAF–PDO and TCF–PDO assembloids, we found that under the treatment-naïve and ertolinib-treatment conditions these two different assembloids demonstrate distinct cytoarchitecture. Moreover, we showed that distinct colocalisation from the TAF–PDO and TCF–PDO assembloid models were recapitulated in different histologic growths patterns in clinical samples. Of note, acinar-specific spatial features were predominantly recapitulated in TCF–PDO assembloids, and solid-specific features in TAFs–PDOs assembloids. This finding highlights the relevance of in vitro patient-derived models to address clinically-relevant questions on the spatial biology of the tumour microenvironment. Moreover, it demonstrates that various in vitro models may be necessary to fully recapitulate specific spatial features from heterogenous disease and provides insights into choosing appropriate systems for downstream functional studies.

When we generated a composite colocatome across the assembloid conditions, we showed that we could reproduce known findings. For example, mesenchymal cancer cells expressing vimentin are generally found in proximity to myofibroblasts, and vice-versa, a known co-occurrence^24^. Through our composite colocatome analysis, we also identified previously unreported cancer–fibroblast spatial features. For instance, we observed that cancer cells and fibroblasts tend to be segregated, rather than being intermixed with each other. Moreover, we found that cancer cells expressing MUC1 (PanCK+Vim-MUC1+, #3) were unlikely to be found in proximity with any fibroblasts. MUC1 is a glycoprotein that provides lubrication and protection to epithelial cells. Its aberrant expression has been involved in cancer development, invasion, metastasis and treatment resistance, including erlotinib^17,25^. Interestingly, both TAFs and TCFs significantly increased MUC1+ cancer cells in the assembloids compared to PDO monocultures, but these cancer cells were not found in the vicinity of any fibroblasts. These results suggests that secreted factors are likely involved in these transformations, as opposed to direct cell–cell interactions.

Given that little is known about the effects of drugs on cytoarchitecture, we studied the effect of erlotinib on TAF–PDO and TCF–PDO assembloids on both the cell composition and spatial organisation. Our findings demonstrate that both TAFs and TCFs can protect cancer cells from erlotinib through unique reorganisation of the tumour–stroma spatial cytoarchitecture rather than promoting heterogeneity or cell subpopulation selection. In this analysis, we observed an enrichment of positive heterotypic colocalisations predominantly involving CAV1-expressing fibroblasts after erlotinib. Several reports, including results from our our group in which we computationally reconstructed the LUAD interactome^26^, concluded on CAV1 being associated with poor outcome in LUAD, while others have suggested the opposite^27^. Additional studies support a high expression of CAV1 in normal lung tissue compared to lung tumours^28–30^. In this case, the colocatome raises further questions regarding whether erlotinib could potentially restore a normal-like lung cytoarchitecture or if the association of CAV1 with a poor outcome may be dependent on specific tumour–stroma interactions.

Further investigation is required to confirm these hypotheses. In addition, we showed that erlotinib-resistant spatial features were mainly recapitulated in TAF–PDO, as opposed to TCF–PDO assembloids. This results align with our previous work regarding the more tumour-promoting role of TAFs versus TCFs^23^ by associating TAF–PDOs’ resistant spatial features with the worse prognostic growth patterns (solid). These results motivate the use of three-dimensional in vitro patient-derived models to identify spatial biomarkers associated with treatment resistance in future functional studies.

Even though the assembloids successfully recapitulated most of the spatial features measured in the LUAD specimens, we acknowledge that a broader spectrum of cell subpopulations were observed in vitro compared to LUAD samples. This may be related to the lower cell density in assembloids, which can affect cell plasticity and phenotype. Although the gross organisation of the cell subpopulations in the tissue samples was maintained in vitro, the in vitro models did not replicate the exact structure of tissues. Future studies should focus on optimising density and account for the other cell compartments in the tumour microenvironment such as immune and endothelial cells. Finally, we recommend expanding this approach to study patient-derived models bearing a range of cancer mutations to assess the spatial reorganisation following treatment as a function of underlying mutations.

In conclusion, the colocatome can serve as a spatial readout to characterise and compare the cytoarchitecture from in vitro patient-derived models and clinical samples. These findings presented on the tumour–stroma architecture of LUAD provide insights into known and novel spatial configurations between cancer cells and fibroblasts in the TME and their potential implications in therapeutic resistance. The colocatome analysis can be applied more broadly and serve as a roadmap to characterise and eventually alter the cytoarchitecture of any complex tissue.

## Methods

### Assembloid cultures

The patient-derived organoids (PDOs) originated from LUAD fresh specimens were kindly provided by Dr. Calvin Kuo, Stanford University and were grown in DMEM:F12 (1:1) media supplemented with 20 ng/ml human FGF-basic, 1X N-2 supplement, 1X B-27 supplement, 10 µM rock inhibitor, 50 ng/ml human EGF and 1X Normocin at 5% CO^2^ and 37°C in a Cultrex reduced growth factor basement membrane extract solid matrix. The LUAD primary fibroblasts were established by our group as previously described^23^ and grown in RPMI-1640 with L-Glutamine, supplemented with 10% fetal bovine serum (FBS), and 5% antibiotic solution (penicillin/streptomycin), at 5% CO^2^ and 37°C. The cell lines, PDOs and cultures used in this study were not tested for mycoplasma contamination or authenticated. Clinical annotations and histopathological informations can be found in Supplementary Table 3.

PDOs and regionally distinct fibroblasts from the tumour edge (TAFs) and core (TCFs) were used to generate the assembloids. Briefly, confluent PDOs grown in a 3D solid dome of Cultrex matrix were rinsed with PBS 1X, and detached from the bottom of the plate with a tip. Then, the domes containing the PDOs were collected in a tube and incubated with TRYPLE at 37°C for 15 minutes under agitation. After the incubation, the domes were gently dissociated using repeated pipetting. In parallel, primary fibroblasts cultures at passage 2^23^ were grown in a monolayer until reaching 75-80% confluency. Then, cells were rinsed with PBS 1X, incubated 5 minutes with TRYPLE, collected and spun down to remove debris. A 1:1 fibroblast:PDO mixture totalising 300k cells was prepared, spun down to remove the remaining liquid, then resuspended in cold liquid Cultrex. Next, 40 µl of the cell mixture was added at the bottom of a 24-well plate pre-warmed at 37°C. Cell mixtures were incubated at 37°C for up to 30 minutes or until the domes solidified. Then, DMEM:F12 (1:1) media formulation supplemented as described above was added to the wells to cover the solidified assembloids (fibroblasts–PDOs). The Assembloids were cultivated for a period of 7 days, followed by treatment with a 2 µM erlotinib solution, which was replenished every 24 hours. The LUAD PDOs employed in this study carry an EGFR del19 mutation that is responsive to erlotinib. Naïve assembloids did not receive any treatment and were used as controls. After 72h, the assembloids were rinsed with PBS 1X and embedded in OCT. OCT blocks were kept at −80°C until sectioning at 7 µm with a cryostat on coverslips (22 × 22 mm) pre-treated with poly-L-lysine overnight, washed 5x with double-distilled water (ddH2O), and dried.

### Human Studies

Clinical aspects of this study were approved by the Stanford Institutional Review Board (IRB) in accordance with the Declaration of Helsinki guidelines for the ethical conduct of research. All patients involved provided a written informed consent. Collection and use of human tissues were approved and in compliance with data protection regulations regarding patient confidentiality (IRB protocol #15166). Following surgical resection of primary tumours, lung adenocarcinoma specimens were immediately processed to establish primary cell cultures as previously described^23^.

### Immunofluorescence

TAF–PDO and TCF–PDO assembloids sections on coverslips were rinsed 3x with a cold solution of 5% BSA (diluted in PBS 1x) and cells were fixed for 10 minutes in PFA 4% (diluted in PBS 1x). Then, the coverslips were incubated in 0.1% Triton X-100 for 10 minutes, rinsed with PBS 1x, blocked with 10% goat serum diluted in PBS 1x for 30 minutes and incubated with the panCK and vimentin antibodies (see Key Resources Table) in blocking solution in a humidified chamber overnight at 4°C. After 3 washes in PBS 1x, the coverslips were incubated with the Cy3-conjugated goat anti-mouse IgG and Cy5-conjugated goat anti-rabbit IgG in blocking solution for 1 hour at RT in the dark. The coverslips were intensively washed with PBS 1x and mounted on slides with a drop of mounting medium containing DAPI. The sections were observed under a BZ-X800 fluorescence microscope.

### PhenoCycler® image acquisition and data processing

The antibody conjugation and the staining were done according to the manufacturer’s protocol (Akoya Biosciences). Details regarding primary antibody, vendors, catalog numbers and barcode assignments are listed in the Key Resource Table. All imaging acquisition were performed with a PhenoCycler® connected to a Keyence BZ-X800 fluorescence microscope equipped with a 20x objective (Nikon CFI Plan Apo 20x/0.75). Before imaging, the region and the z-stack were configurated on the Keyence software and the number of cycles with targets were designated in the Akoya’s PhenoCycler® Instrument Manager (version 1.29.3.6) as described in the CODEX user manual (Akoya Biosciences, 2021a). Acquired images were processed, stitched, and segmented using the CODEX Processor set at default manufacturer values with a radius of 10 for segmentation (Akoya Biosciences, 2021b). Final data were then viewed using the CODEX Multiplex Analysis Viewer (MAV) plugin (Akoya Biosciences, 2021c) for FIJI (ImageJ)^31^ and the EnableMedicine Visualizer tool (beta version).

### Cell assignment with CELESTA

CELESTA is an an unsupervised machine learning algorithm for individual cell identification using the cell’s marker expression profile and, when needed, its spatial information^11^. The CELESTA package was downloaded from GitHub under https://github.com/plevritis-lab/CELESTA. As a first step, a cell type signature matrix that relies on prior knowledge of markers was designed and used as CELESTA input, along with the segmentation data as previously described^11^. Our initial cell type signature matrix included all the possible combinations of a pre-selected set of markers from our antibody panel to identify cancer cells (panCK, VIM, MUC1) and fibroblasts (αSMA, CD90, PDGFRb, PDPN, CAV1), along with other general markers to identify subpopulations. Every cell subpopulation (designated by unique combination of markers) with a cell fraction smaller than 2% across all samples were removed from the cell type signature matrix before a subsequent iteration of CELESTA, until reaching a final cell type assignment. Note that Vim+PanCK-EpCAM-cells were assumed as fibroblasts, and that we used the expression of EpCAM to confirm that PanCK- cancer cells were not fibroblasts. The cell type signature matrix and the CELESTA thresholds can be found in Supplementary Table1.

### Colocatome analysis

To quantify the tumour–stroma cytoarchitecture within the assembloids, we used the colocation quotient and identified positive (proximate) and negative (distant) cell–cell spatial features between each cell subpopulation as previously described^11,12^. Briefly, we use the colocation quotient to quantify how a cell subpopulation colocates spatially with another cell subpopulation among a set of nearest neighbours, defined here as 20. We calculated the colocation quotient for the pairwise cell types identified under naïve and treatment conditions using the following equation: CLQ_b→a_ = (C_b→a_/N_a_) / (N_b_/(N− 1)) where C is the number of cells of cell type b among the defined nearest neighbors of cell type a, N is the total number of cells and N_a_ and N_b_ are the numbers of cells for cell type a and cell type b. We use a Euclidean distance bandwidth parameter, which is determined based on the segmented X and Y coordinates, with a default value set to 100. We assessed the significance of the CLQ values obtained by randomly permuting 500 times the cell labels while preserving the subpopulation proportions. CLQ values falling outside or at the tail of the distribution generated by the permutation analysis were considered significant, whereas values within the distribution were deemed non-significant, as they can be reproduced after spatial randomisation. Percentile values < 0.05 or > 0.95 were considered as significant. The CLQ values were nomalised based on the mean and range of values derived from the permutation analysis. This process also considered the number of cells within each subpopulation. Subpopulations with a low cell count were more likely to yield a broader distribution of CLQ values during the permutation analysis. This broader distribution resulted from the substantial impact of random label sampling on CLQ value calculations. A wider distribution increases the likelihood of categorising the original CLQ value as statistically insignificant. The normalisation achieved through the permutation analysis facilitates not only spatial feature comparisons but also enables the comparison of different conditions from the same, or independent experiements. CLQs were normalised according to the following formula: (Observed CLQ - Mean CLQ)/(Max CLQ – Mean CLQ).

CLQ calculations were repeated in the periphery and centre regions only of the same assembloid to evaluate whether specific spatial features were enriched in a particular region. To determine each region, the assembloid sections were considered as circular, and periphery and centre zones were established with a consistent ratio to the diameter, ensuring uniform zone ratios regardless of the total area that can vary between samples (Extended Data Fig. 3).

### Composite colocatome of assembloids

To generate our composite colocatome using both assembloid models and treatment conditions, we identified colocalisations consistent across assembloid biological replicates for each condition (TAF, TCF, naïve and erlotinib-treated assembloids). Briefly, all negative colocalisations (cells repelling each other or segragated) were assigned to −1, and positive colocalisations (cells found in proximity) were assigned to 1. Unsignificant spatial features were assigned to 0. Cell pairs colocalised under one condition, but negatively colocalised under another were assigned to 1, as the reference colocatome intented to generate a matrix of possibilities of cells that can be found in proximity in some, but not necessarily all instances. The statistically significant spatial features were combined to establish a comprehensive reference colocatome between cancer cells and fibroblast subpopulations from the LUAD. Next, we used hierarchical clustering on the full (homo- and heterotypic spatial features) or partial (heterotypic spatial features) colocatome to group cell subpopulations into subcommunities with similar colocalisation partners. Then, we ordered the subcommunities according to their likelihood of being found in proximity or segragated from each other (clusters), which led to a precise characterisation of the cytoarchitecture.

### LUAD Clinical Samples

To validate the clinical significance of our assembloid-derived colocatome, we transposed the cell subpopulations identified in the assembloids to a small LUAD cohort available at https://www.enablemedicine.com/XX12345678XX (accession number to come at the time of publication). The cell signature matrix developed for the assembloids was used to guide the identification of fibroblast and cancer cell subpopulations in LUAD specimens (Supplementary Table 3) of unknown cell composition and architecture. Lepidic, acinar and solid regions from a subset of samples were defined by a pathologist, overlayed on the Phenocycler images using the EnableMedicine Beta visualiser tool, and cells were assigned to a histological growth pattern (lepidic, acinar or solid). The LUAD samples were then broken into histopathological regions associated with a growth pattern, on which we repeated the analysis pipeline described above. Significant CLQ values consistent across more than half of the number of regions of a same growth pattern were considered as significant spatial features to reconstruct the cancer-fibroblast lepidic, acinar and solid colocatomes. Spatial features that resisted or emerged after erlotinib treatment were carried out to LUAD specimens. Briefly, resistant spatial features that overlaped between assembloids and lepidic, acinar or solid growth patterns were used to identify spatial biomarkers associated with resistance on clinical specimens.

### Statistical analysis

Statistical testing for the CELESTA cell proportions were performed with GraphPAD Prism Software 9.5.1 using two-way ANOVA followed by multiple comparisons. Cell density analyses and cell fractions in assembloids before and after erlotinib treatment were performed using one-way ANOVA, followed by multiple comparisons. Error bars in the figures represent the SD. Colocation cluster overlaps with histopathological regions were calculated using a hypergeometric test. P values < 0.05 were considered as significant.

## Supporting information

Key Resource Table

## Data availability

Assembloid imaging data are hosted at EnableMedicine (accession number will be available at the time of publication). Segmentation data will be provided upon reasonable request for research purposes. Reagents and antibodies used for this study are available in the Key Resource Table and Reporting Summary. Further information and requests for resources and reagents should be directed to, and will be fulfilled by, the lead contact, S.K.P.. This study did not generate new unique reagents or cell lines.

## Code availability

The R scripts used in this study will be deposited on the Plevritis lab Github at https://github.com/plevritis-lab/ at the time of publication.

## Acknowledgements

The authors thank Enable Medicine for providing visualisation and analytical tools, as well as technical support for mIF images.

## Author contributions

Study Conception & Design: G.B. and S.K.P.; Performed Experiment and Data Collection: G.B., I.L., A.B. and Y.L.; Data Analysis: G.B., W.Z., I.L, I.I., S.K.P.; Interpretation of data analysis: G.B., W.Z. I.L., I.I, L.T, W.T., J.B.S., A. J.G. and S.K.P.; Writing the first draft: G.B; Figures Design: G.B; Patient Sample Management : A.B., C.K., J.B.S. and W.T.; Supervision: J.B.S., A.JG. and S.K.P. All authors contributed to manuscript editing and revision.

## Supplementary methods

### Flow Cytometry Analysis

After TRYPLE incubation, PDOs were dissociated into single-cell suspension. Cells were counted and aliquots of 1×10^6^ cells per condition were prepared. The samples were incubated for 5 minutes with 1 µL of Zombie Aqua™ fixable viability dye in PBS 1X, then washed with flow cytometry buffer (FCB, 0.5% BSA, 0.02% NaN_3_ and 2mM EDTA in PBS 1X) and centrifuged (500 × g, 5 minutes) before adding PFA at a final concentration of 1.6% for 10 minutes at room temperature in FCB. Cells were then centrifuged at 500 × g for 5 minutes at 4°C to pellet cells, PFA was removed, and cells were washed again with FCB. Cells were either long-term stored at −80°C in 500 µL of FCB or permeabilised with 100 µL of eBioscience™ Permeabilisation Buffer diluted at 1X concentration for 30 minutes on ice with a master mix of primary antibodies (see Key Resource Table). After the incubation with primary antibodies, cells were washed with FCB and spun down at 500 × g for 5 minutes at 4°C (2X). Then, cells were resuspended in 500 μl of FCB, strained and analysed using a BD LSRFortessa™ X-20. Results were analysed using Cytobank single-cell analysis software with the gating strategies described in Extended Data Fig. 6.

### EMT analysis with PhenoSTAMP

PHENOSTAMP was downloaded from GitHub under https://github.com/anchangben/PHENOSTAMP. Briefly, FCS files previously gated in Cytobank according to singlets and Live/Dead (Extended Data Fig. 6), were uploaded into R, and the PHENOSTAMP algorithm was used to project the PDOs on the 2D EMT–MET state map, as previously described here^1^.

---

**Extended Data Fig. 1.**
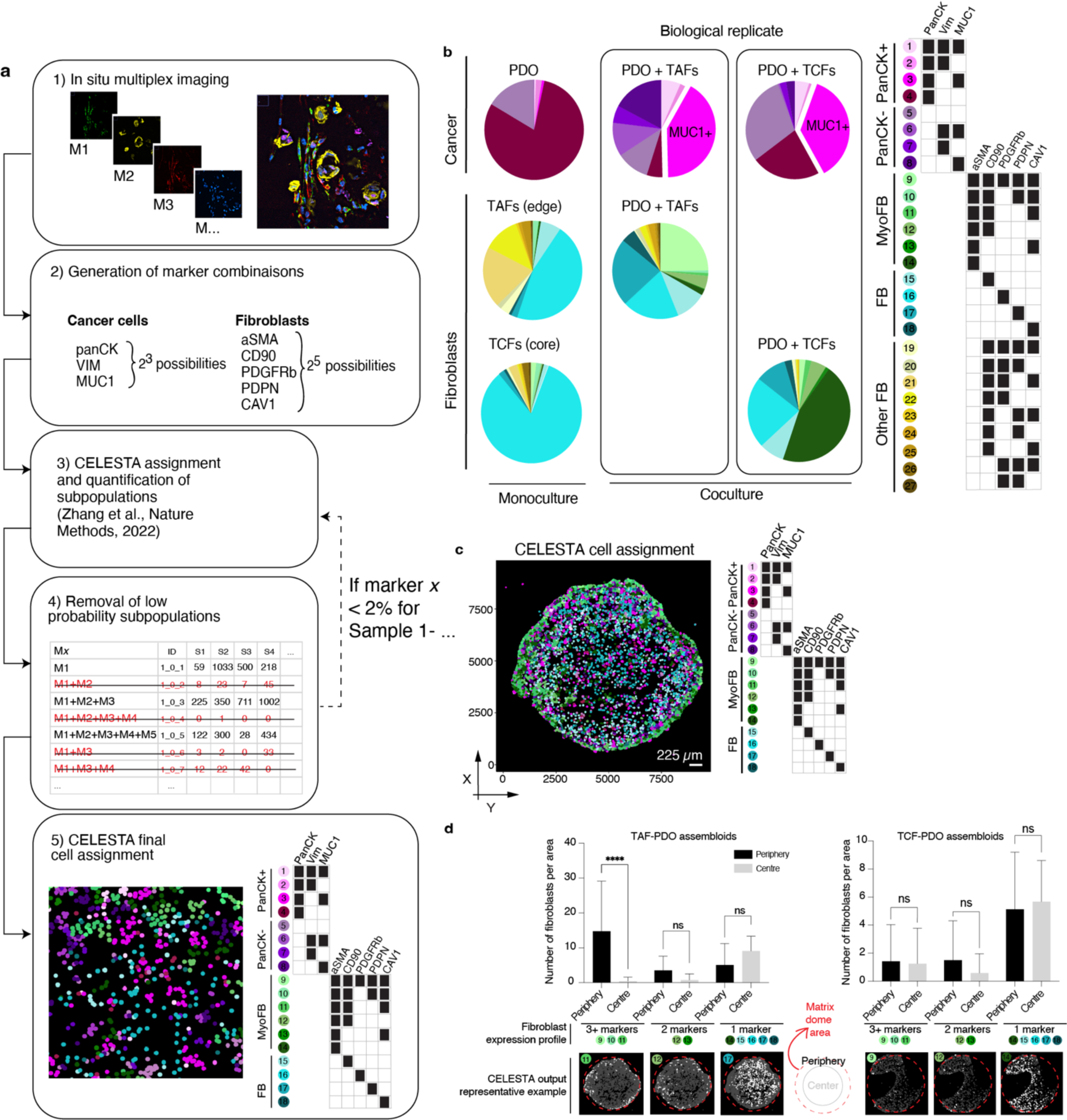
Cell heterogeneity in LUAD assembloids. **a,** Study workflow for subpopulation identification using the iterative nature of CELESTA. **b,** Representative example of subpopulation proportions in TAF–PDO and TCF–PDO assembloids (TAFs from LUAD specimen 1 and TCFs from LUAD specimen 2). **c**, Representative image of assembloid shown as a dot plot where each dot represents a cell, located at the cell centroid, and is color-coded by its cell subpopulation identified with CELESTA. **d**, Bar graphs representing the number of fibroblasts per area (1000 x 1000 a.u.) in each zone of assembloid, namely the periphery versus centre. Quantification, out of six representative areas per region (centre, periphery) from two independent biological replicate per condition. Statistical significance was determined using one-way ANOVA. Error bars indicate standard deviation of the mean.

**Extended Data Fig. 2.**
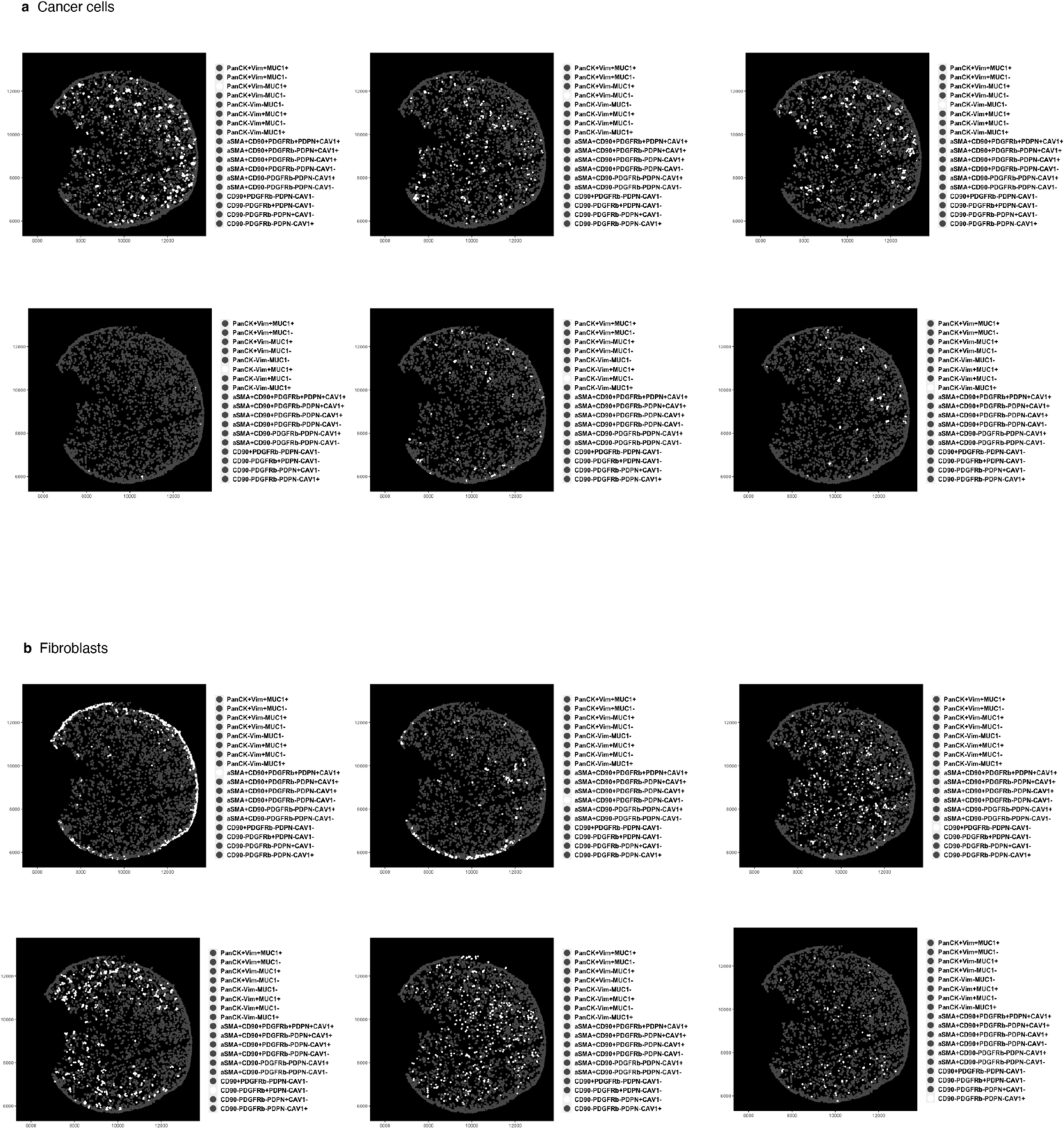
TAF-PDO assembloid from LUAD specimen #1 showing dot plots where each dot represents a cell, located at the cell centroid, and is white color-coded by **(a)** cancer cell and **(b)** subpopulation identified with CELESTA.

**Extended Data Fig. 3.**
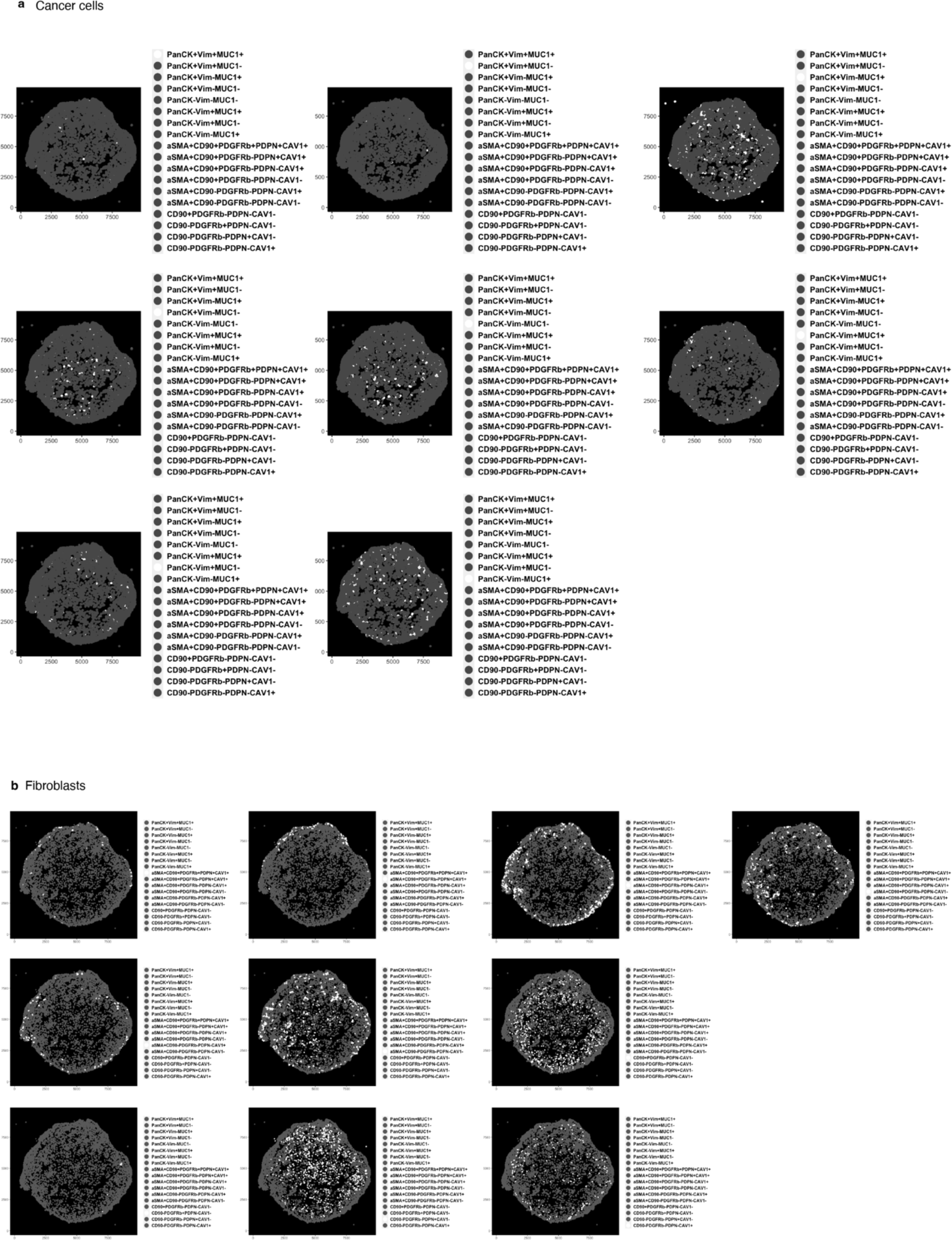
TAF-PDO assembloid from LUAD specimen #2 showing dot plots where each dot represents a cell, located at the cell centroid, and is white color-coded by **(a)** cancer cell and **(b)** subpopulation identified with CELESTA.

**Extended Data Fig. 4.**
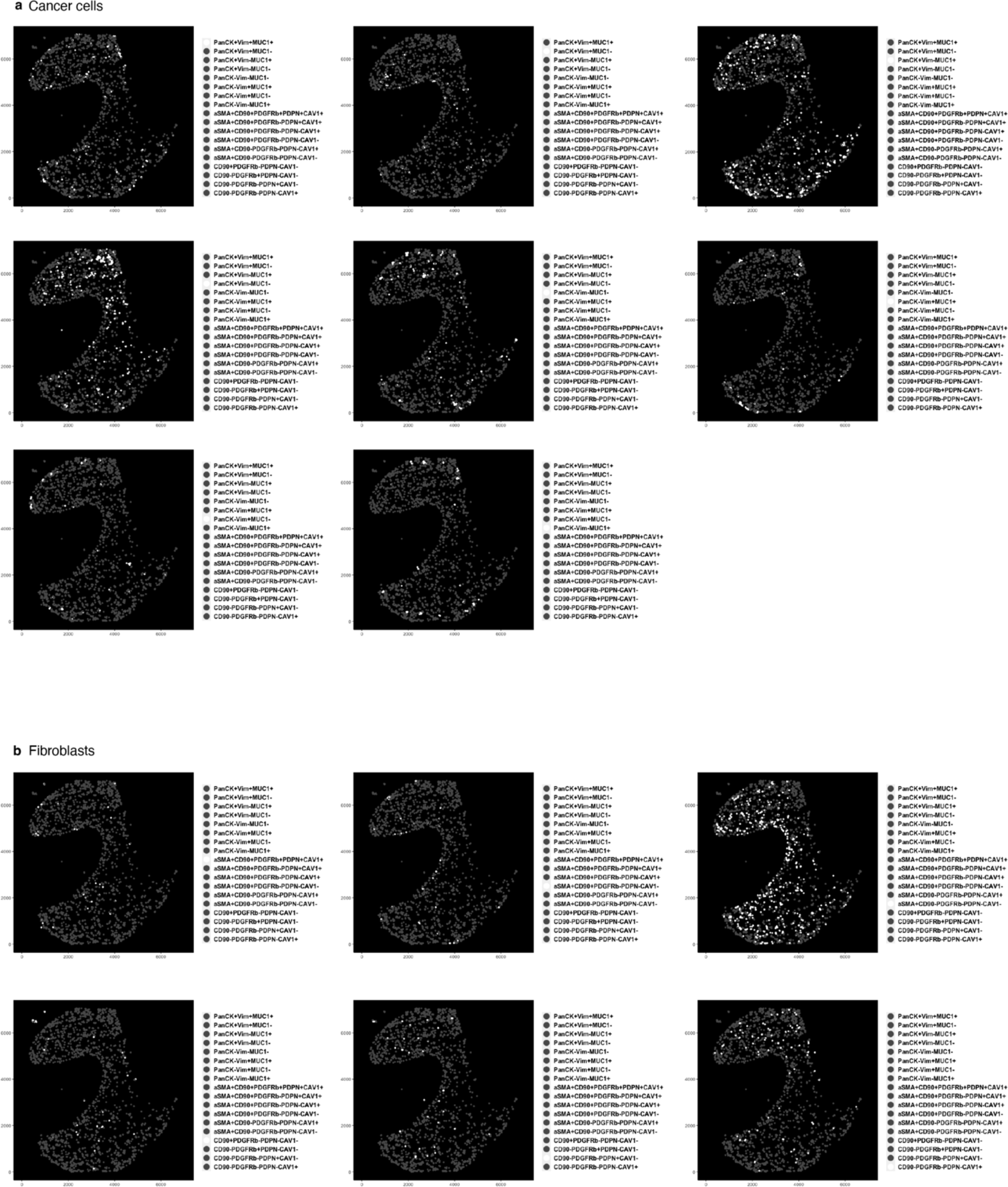
TCF-PDO assembloid from LUAD specimen #1 showing dot plots where each dot represents a cell, located at the cell centroid, and is white color-coded by **(a)** cancer cell and **(b)** subpopulation identified with CELESTA.

**Extended Data Fig. 5.**
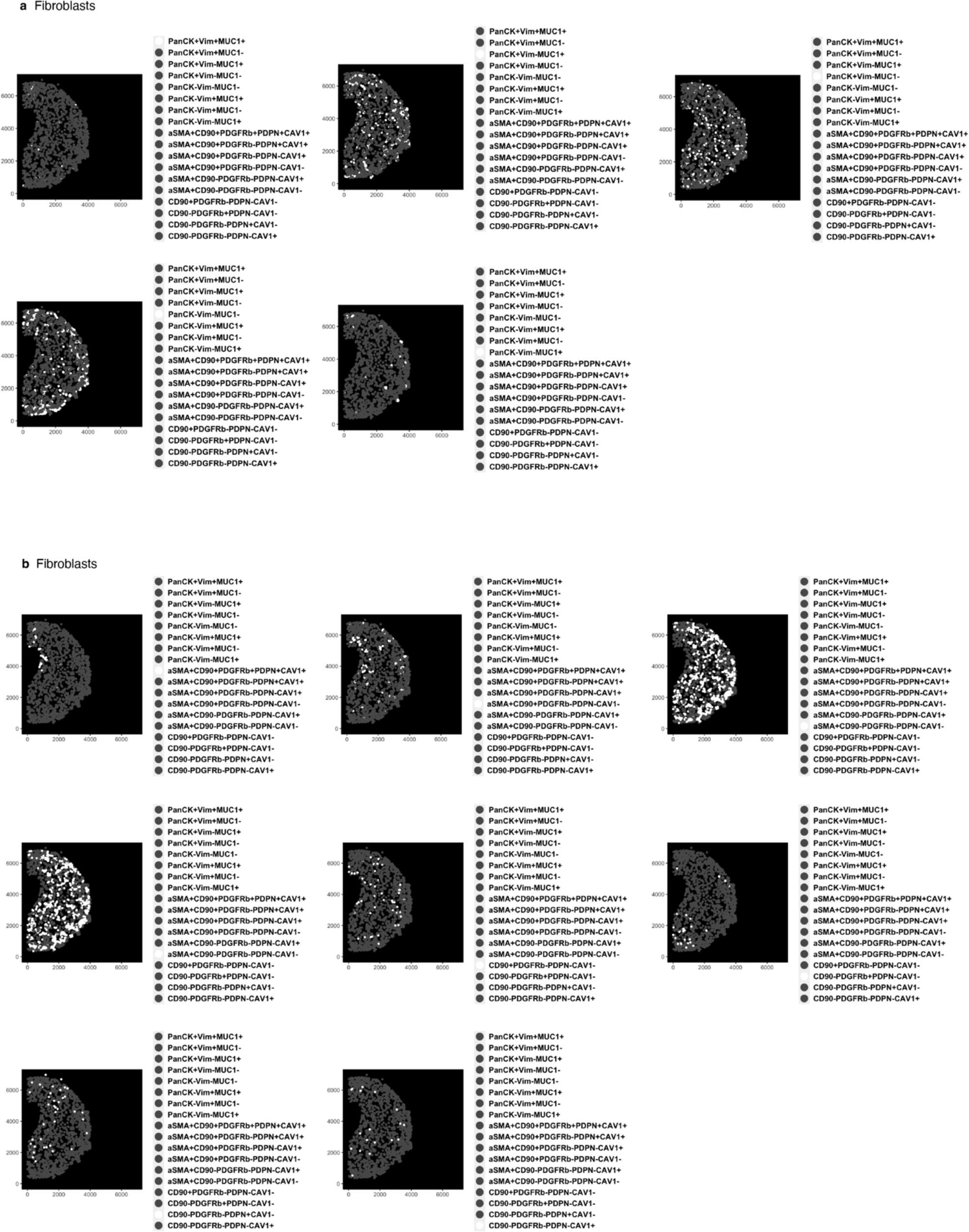
TCF-PDO assembloid from LUAD specimen #2 showing dot plots where each dot represents a cell, located at the cell centroid, and is white color-coded by **(a)** cancer cell and **(b)** subpopulation identified with CELESTA.

**Extended Data Fig. 6.**
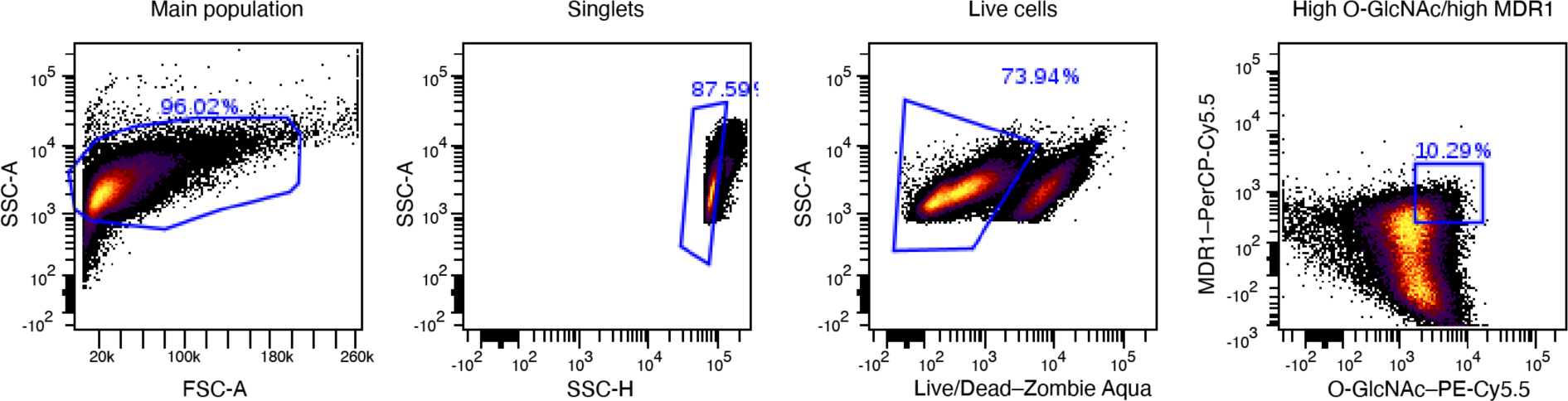
Flow cytometry gating strategy. Gating strategy used for the PHENOSTAMP projections of PDOs analysed with flow cytometry over the erlotinib time course.

**Extended Data Fig. 7.**
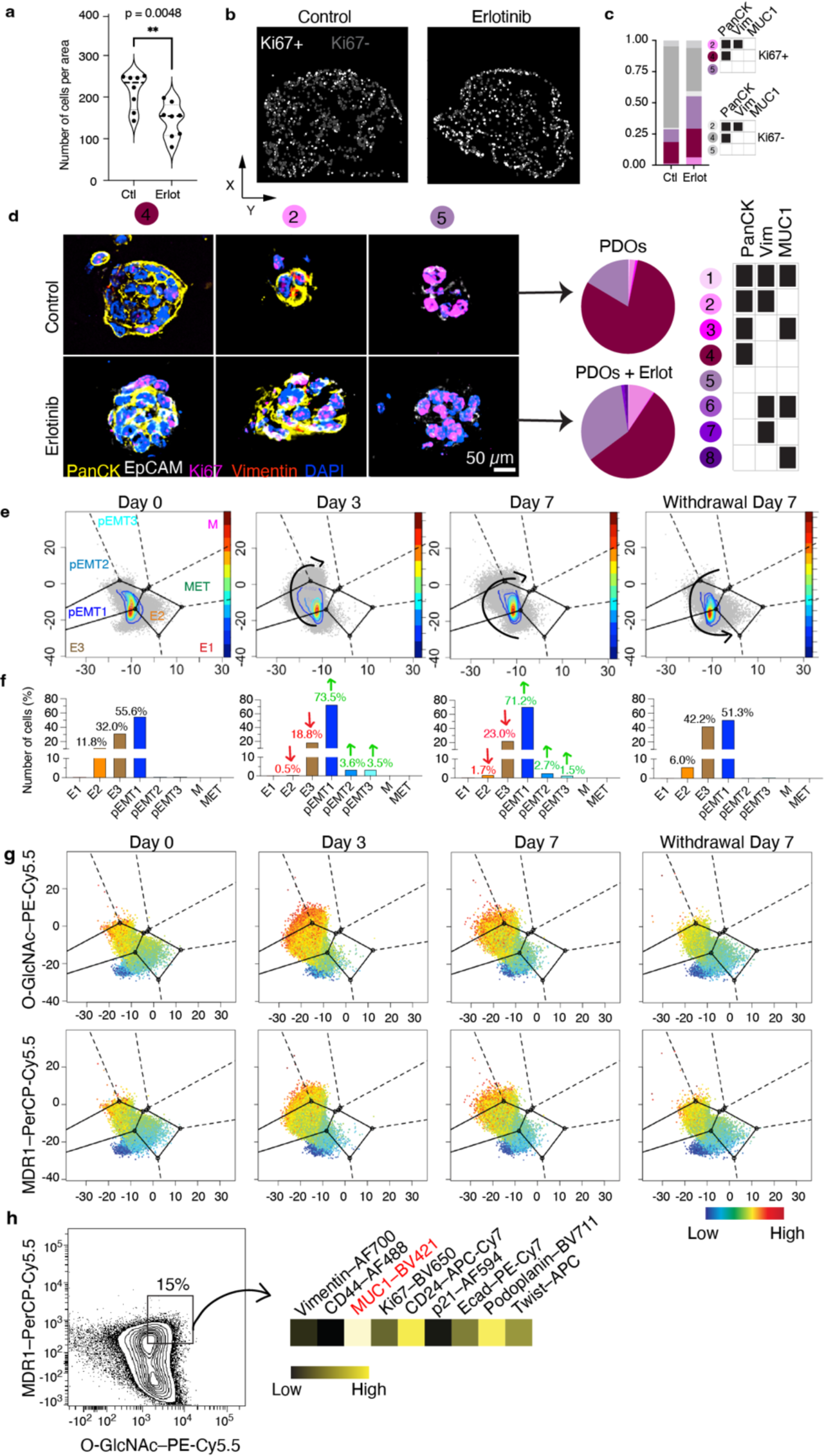
Effect of erlotinib on LUAD PDOs. **a**, Violin plot representing cell density. Quantification out of 8 representative areas per condition. Statistical significance was determined using two-tailed Student’s t-test. **b,** Dot plot where each dot represents a cell, located at the cell centroid showing proliferative (Ki67+) and non-proliferative cancer cells. **c**, Bar graph showing the proportions of proliferative cells before and after erlotinib. **d**, Representative PDO images showing the main subpopulations present in monocultures. **e**, PHENOSTAMP projections of PDOs analysed with flow cytometry over erlotinib time course and (**f**) bar graph showing cell quantifications in each EMT state. **g**, PHENOSTAMP projections showing the single-cell expression of known erlotinib resistance markers (O-GlcNAc and MDR1). **h**, Flow cytometry contour plot displayed the expression profile of highMDR1/highO-GlcNAc partially EMT cancer cell subpopulation. Experiments involving monocultures were conducted once as control. E, epithelial; pEMT, partialEMT; M, mesenchymal; T, transition; O-GlcNAc, O-linked β-N-acetylglucosamine; MDR1, multidrug resistance protein 1; CD, cluster of differentiation; Ecad, e-cadherin.

**Extended Data Fig. 8.**
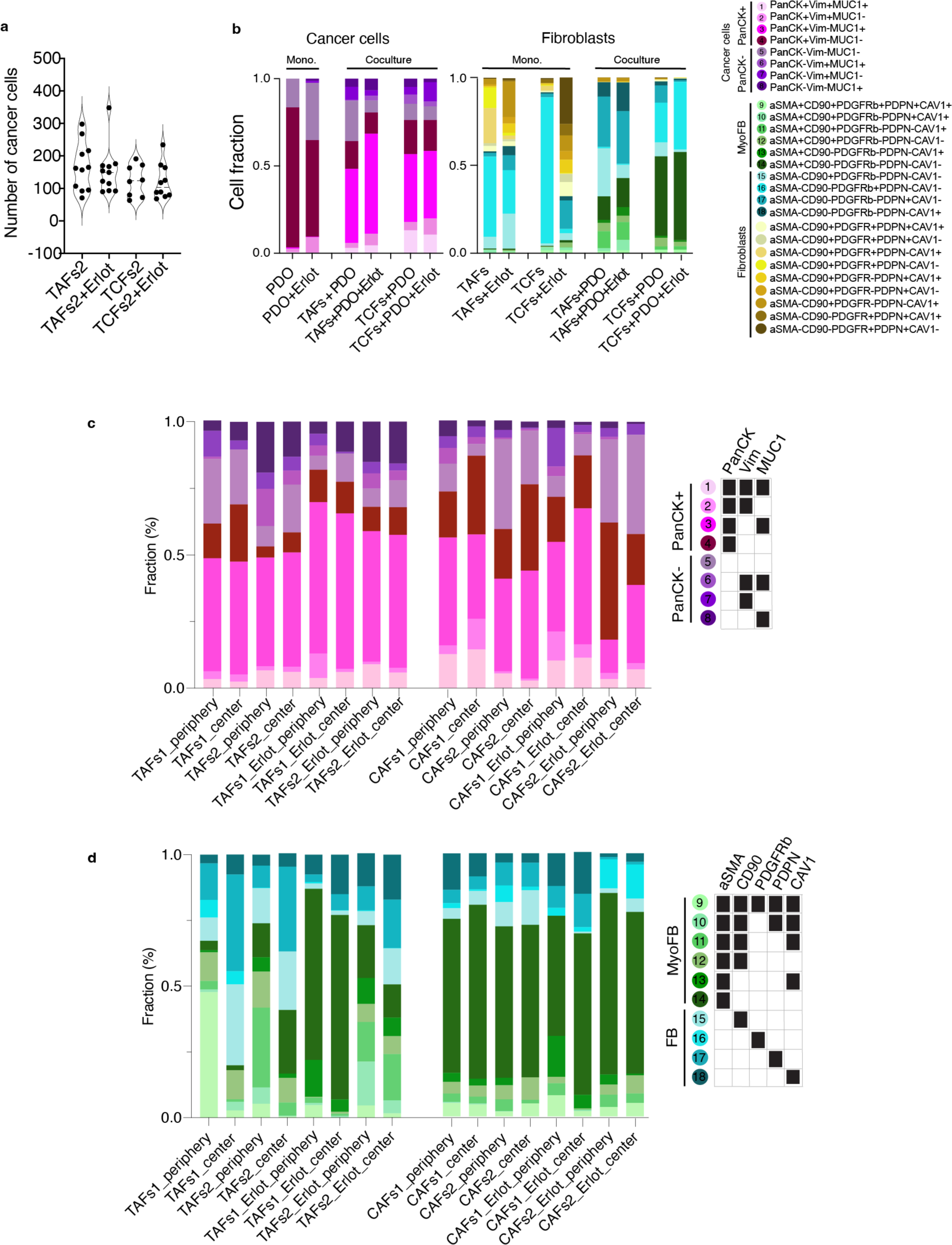
Effect of erlotinib on LUAD assembloids heterogeneity. **a,** Violin plot representing cell density for TAF–PDO specimen #2. Quantification out of 8 representative areas per condition. **b,** fraction of each cell subpopulation before and after erlotinib from two independent biological replicate per condition except monocultures (n = 1). **c,d,** Cell proportions in periphery vs centre assembloid regions.

**Extended Data Fig. 9.**
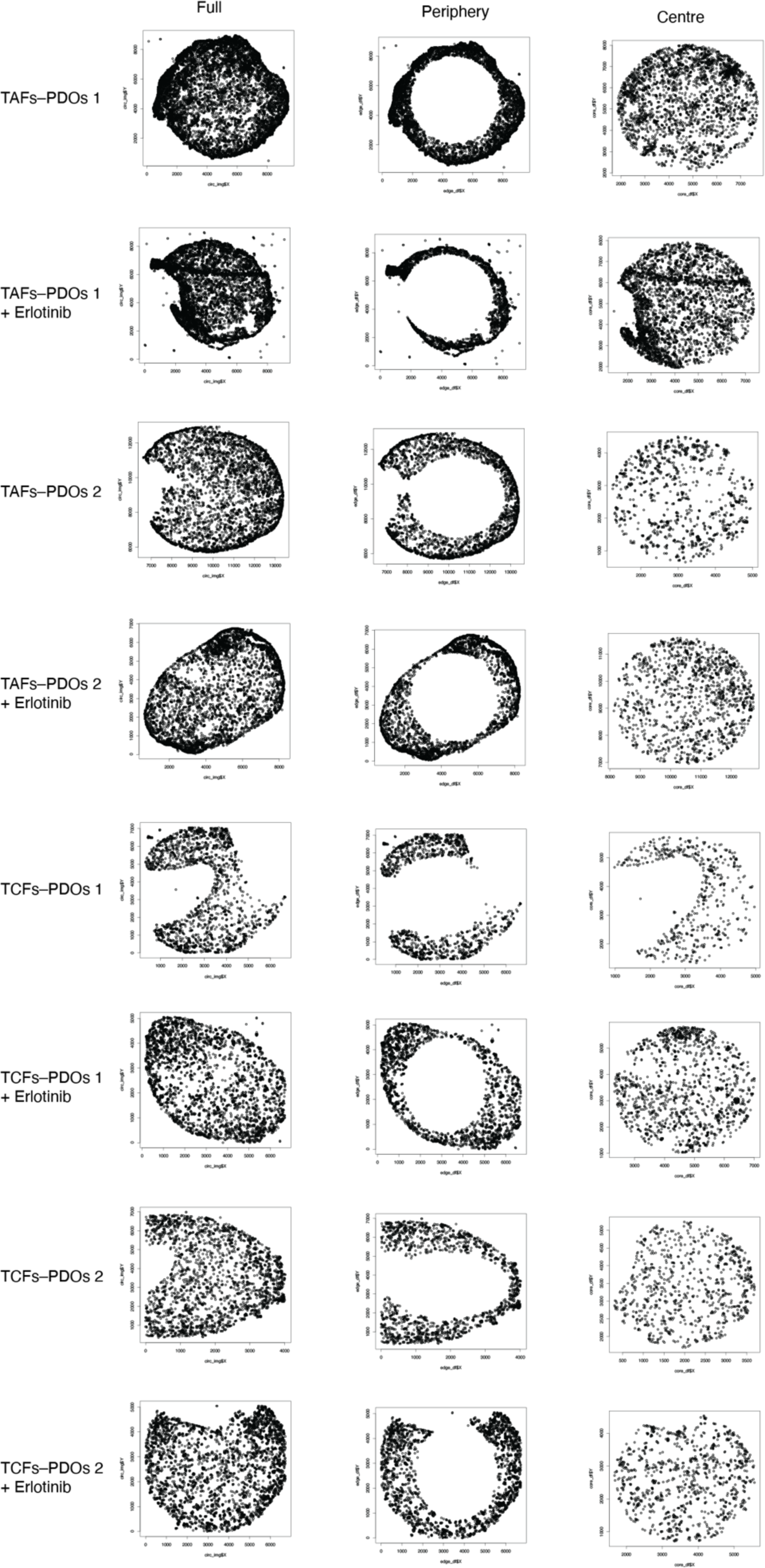
Assembloid data. Dot plots where each dot represents a cell showing the cells included in the analyses using the full, periphery or centre data for each assembloid condition.

**Extended Data Fig. 10.**
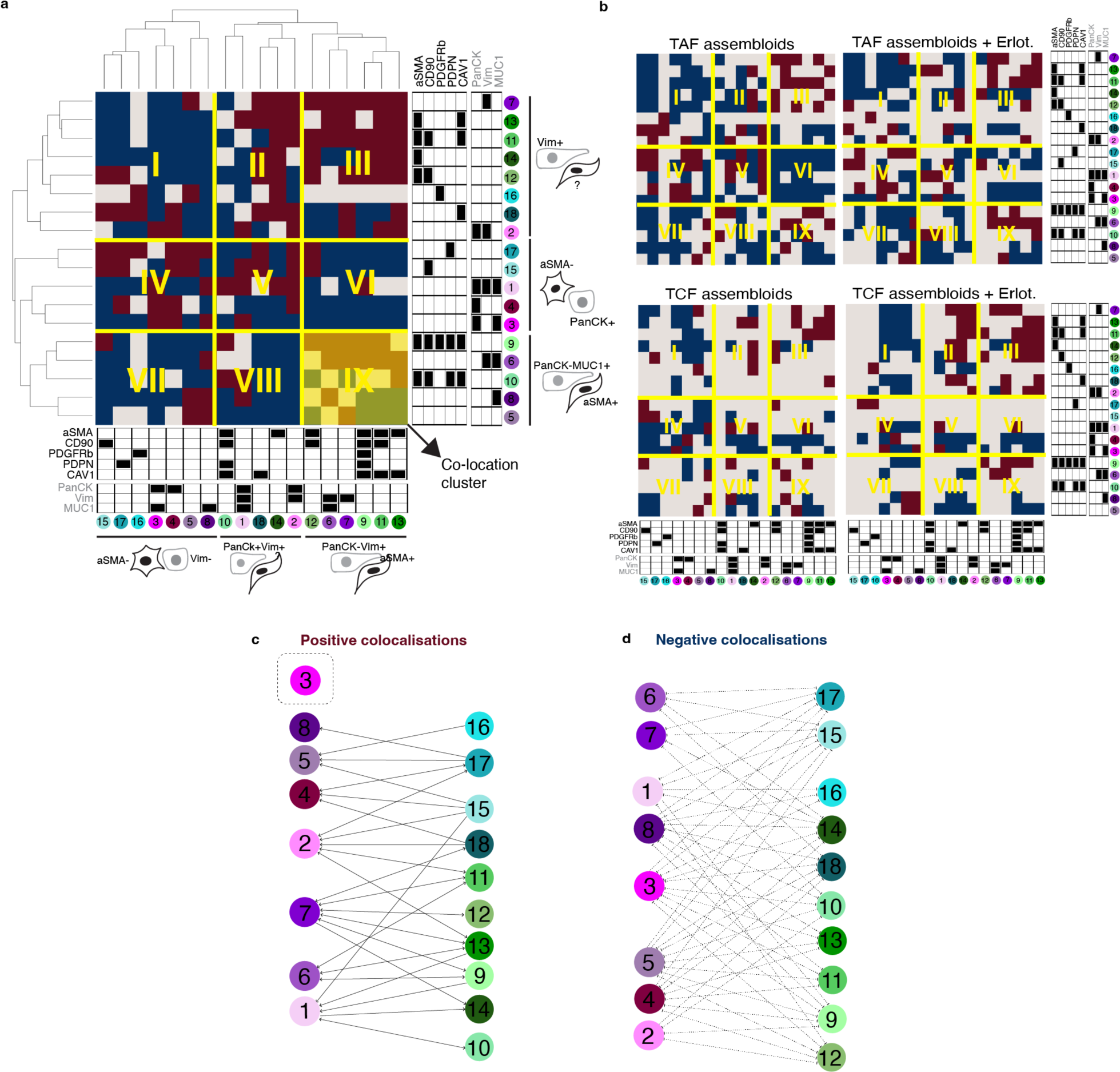
The colocatome of cancer–fibroblast assembloids. Hierarchical clustering of the full (**a**) colocatome and (**b**) partial colocatome per condition. Bipartite plot summarising positive (**c**) and negative (**d**) heterotypic colocalisations.

**Extended Data Fig. 11.**
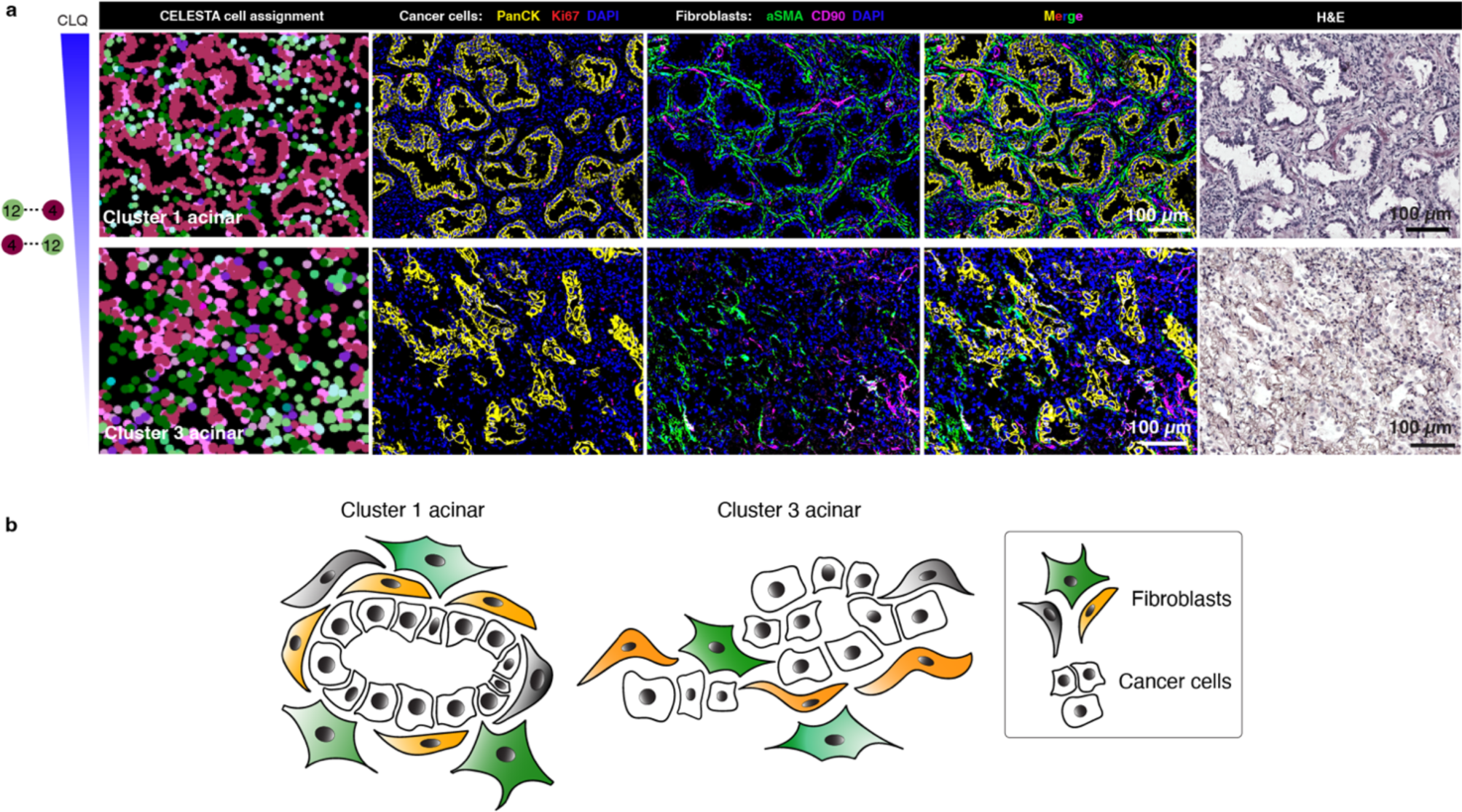
The colocatome influences acinar spatial features. **a,** Representative PhenoCycler images showing cancer cells and fibroblasts in acinar regions from clusters 1 and 3 and (**b**) cartoon summary of their cytoarchitecture.

**Extended Data Fig. 12.**
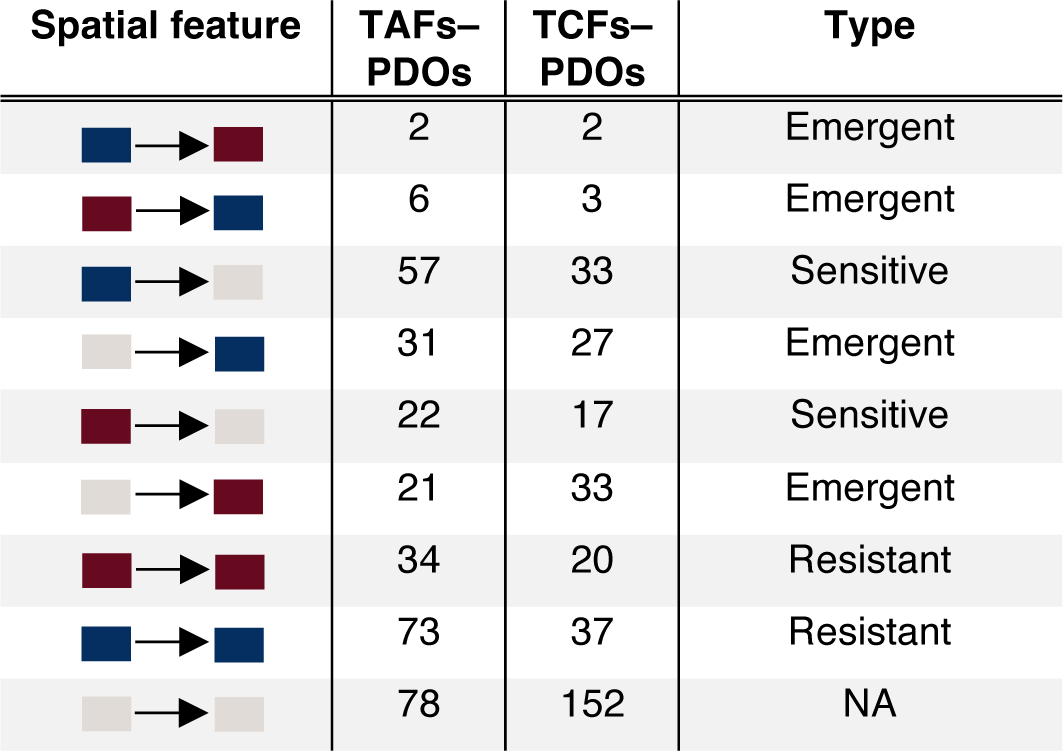
Erlotinib-induced changes in the colocatome. Persistent, emergent, and sensitive colocalisations measured in erlotinib vs naïve assembloids. Blue boxes represent negative colocalisations, red boxes represent positive colocalisations, grey boxes: insignificant colocalisations.

**Extended Data Fig. 13.**
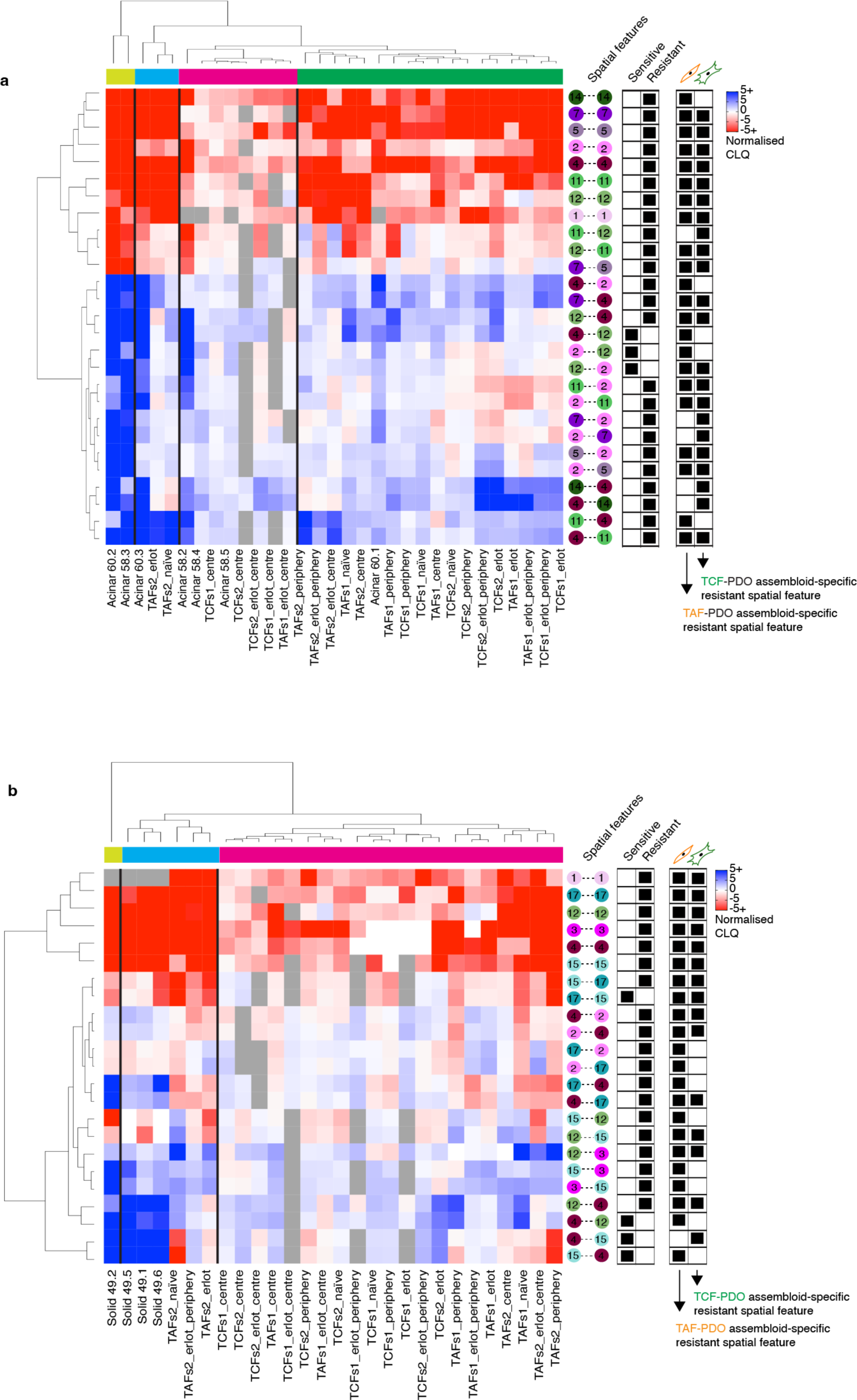
Integrating the colocatome of assembloids with LUAD specimens. Heatmap of the spatial features identified from the LUAD **(a)** acinar and **(b)** solid regions integrated with the erlotinib-resistant and sensitive spatial features inferred from the reference colocatome.

**Extended Data Fig. 14.**
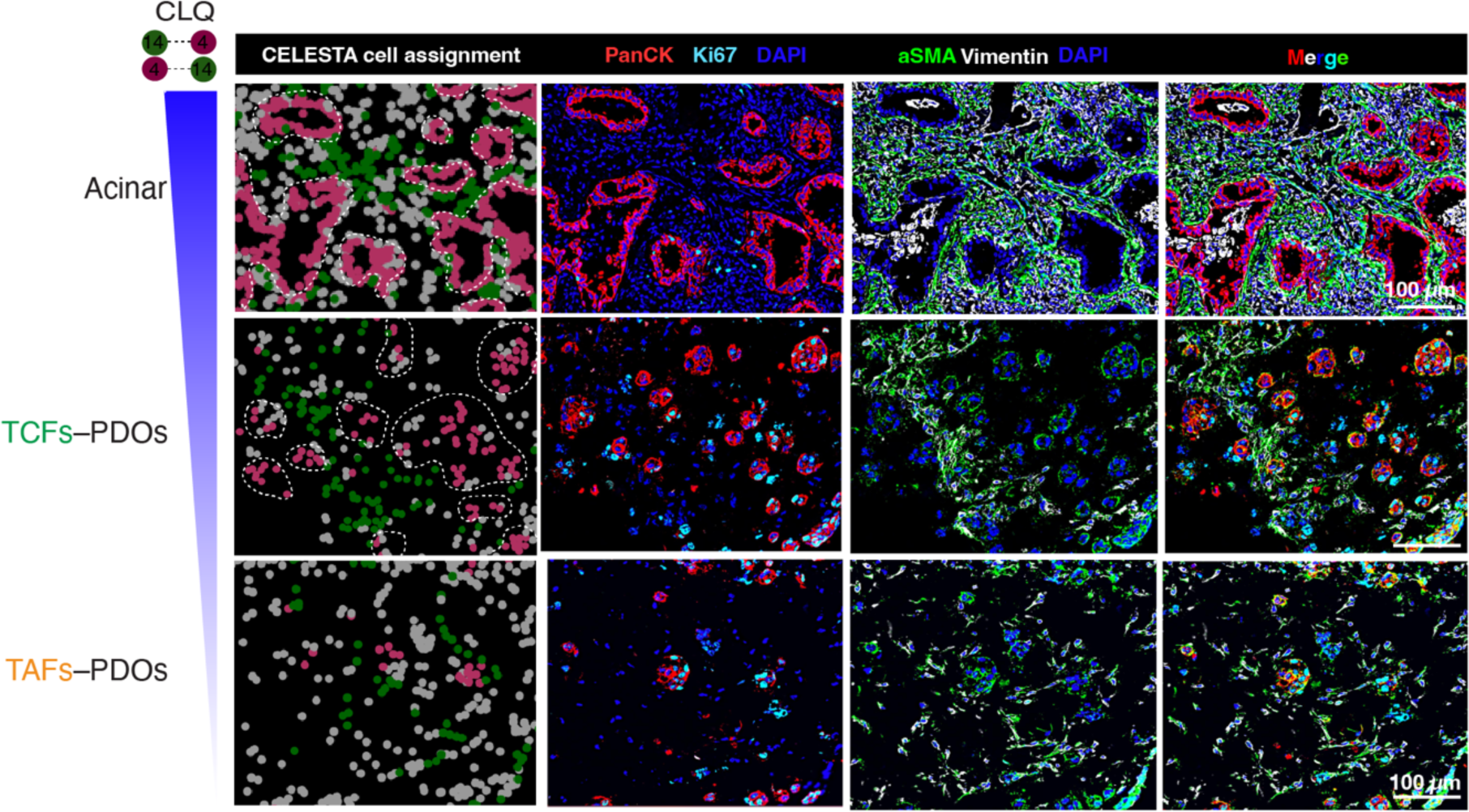
The colocatome influences acinar spatial features. **a,** Representative PhenoCycler images showing cancer cells and fibroblasts in acinar regions from clusters 1 and 3 and (**b**) cartoon summary of their cytoarchitecture. **c**, acinar-like spatial feature recapitulated in TCF–PDO, but not TAF-PDO assembloids.

