## Supplementary table 3 for "The colocatome as a spatial -omic reveals shared microenvironment features between tumour–stroma assembloids and human lung cancer"

| Primary fibroblast cultures | | | | | | | | | | | | | | | | | | |
| --- | --- | --- | --- | --- | --- | --- | --- | --- | --- | --- | --- | --- | --- | --- | --- | --- | --- | --- |
| Specimen | **TAFs** | | **TCFs** | | **Tumour size (cm)** | **Differentiation status** | | **Prior Treatment** | | **Comorbordities** | | **Smoker status** | | **Common mutations** | **Classification** | | **Node status** | |
| 1 | X | | X | | 3.5 | Well | | None | | None | | Never | | EGFR exon 19 deletion | Acinar 85%, lepidic 10%, pipillary 5% | | N0 | |
| 2 | X | | X | | 2.6 | Moderate | | None | | Hypertension | | Former | | None | Papillary 50%, acinary 30%, lepidic 20% | | N1 | |
| Patient-derived Organoids | | | | | | | | | | | | | | | | | | |
| Specimen | **Sex** | **Stage** | | **Race** | | | **Tumour size (cm)** | | **Differentiation status** | | **Prior Treatment** | | **Smoker status** | | **Common mutations** | **Classification** | | **Node status** |
| PDO | M | 4 | | Black or African American | | | >1 mm - < 5mm | | Unknown | | None | | Never | | EGFR exon 19 deletion | Adenocarcinoma (NOS) | | N3 |
| LUAD pathological samples | | | | | | | | | | | | | | | | | | |
| Specimen | **Sex** | **Stage** | | **Comorbordities** | | | **Tumour size (cm)** | | **Differentiation status** | | **Prior Treatment** | | **Smoker status** | | **Common mutations** | **Classification** | | **Node status** |
| Adeno 49 | ? | 3 | | None | | | 4.8 | | Moderate-Poor | | None | | Never | | None | Solid predominant | | N1 |
| ADENO 58 | F | 2a | | Arthritis, diabetes, hypertension, hyperlipidemia, tendinitis | | | 3.9 | | Well | | None | | Never | | EGFR c.2573T>G | Acinar predominant | | N2 |
| ADEnO 60 | M | 2a | | Occupational exposure, depression, pneumothorax | | | ? | | Well | | None | | Former | | EGFR c.2236_2250del, TP53 c.434T>C | Acinar predominant | | N0 |

**Supplementary Table 5.** **Clinical annotations and histopathological information of LUAD specimens and primary cells cultures used in this study.**
